# Stem-like PD-1^+^TCF-1^+^ CD8^+^ T cells result from helpless priming and rely on CD4^+^ T-cell help to complete their cytotoxic effector differentiation

**DOI:** 10.1101/2025.10.18.683204

**Authors:** Julia Busselaar, Douwe M. T. Bosma, Mo D. Staal, Xin Lei, Yanling Xiao, Jannie Borst

## Abstract

**Background:** Antigen-specific CD8^+^ T cells can be in a stem-like PD-1^+^TCF-1^+^ differentiation state that progresses into terminal exhaustion in cancer and chronic infection. These stem-like cells are important, since they are the responders to PD-1 targeted immunotherapy.

**Methods:** We use mouse vaccination and tumor models to delineate the effects of CD4^+^ T-cell help during priming on the differentiation fate of stem-like CD8^+^ T cells.

**Results:** In absence of help signals, stem-like CD8^+^ T cells do not differentiate into effector cytotoxic T cells (CTLs) and accumulate in the draining lymph node (dLN). When help signals are delivered, either during or after priming, stem-like CD8^+^ T cells proliferate and differentiate into circulating effector cells. Stem-like CD8^+^ T cells raised by vaccination in presence or absence of CD4^+^ T-cell help have an identical transcriptome, which they share with stem-like cells defined in mouse models of cancer and chronic infection. Regardless of expression of strong helper epitopes as present in our vaccine, the immunogenic MC38 tumor does not prime helped effector CD8^+^ T cells but primarily stem-like cells.

**Conclusions:** Our data support the proposition that stem-like CD8^+^ T cells are helpless cells that lie at the bifurcation point of CD8^+^ T-cell effector- and exhaustion trajectories. Effective delivery of help signals to stem-like CD8^+^ T cells to drive their expansion and CTL effector differentiation is an important therapeutic challenge in cancer and other conditions that lead to T-cell exhaustion.

**SUMMARY:** *What is already known on this topic:* PD-1^+^TCF-1^+^ stem-like CD8^+^ T cells are the progenitors of exhausted cells in cancer and chronic infection and the main responders to PD-(L)1-targeting cancer immunotherapy. Stem-like CD8^+^ T cells can differentiate towards functional effectors or exhausted cells, but the question remains which signals drive this fate decision during priming.

*What this study adds:* This study shows that PD-1^+^TCF-1^+^ stem-like CD8^+^ T cells are cells that have not (yet) received CD4^+^ T-cell help signals during priming. In a vaccination setting, providing help signals to stem-like CD8^+^ T cells, even after priming, drives their differentiation towards functional CTLs. However, in a tumor context, CTL differentiation is impaired even when helper epitopes are present in the tumor and a prime a CD4^+^ T-cell response.

*How this study might affect research, practice or policy:* This study elucidates the origin of stem-like CD8^+^ T cells, and identifies CD4^+^ T-cell help as critical factor to steer their differentiation in the direction of optimal CTLs. It also highlights that expression of immunogenic helper epitopes by tumor cells does not guarantee that CD4^+^ T-cell help for CTL differentiation is delivered. These findings expand the current CD8^+^ T-cell differentiation model and provide a rationale to optimize immunotherapeutic strategies.

## BACKGROUND

In a cytotoxic T lymphocyte (CTL) response that results in antigen clearance, the differentiation of newly activated CD8^+^ T cells is described as a trajectory ranging from the naïve state, via central memory (T_CM_) and effector memory (T_EM_) precursor states to a terminally differentiated effector CTL state^1,2^. In situations where antigen persists, such as chronic infection and cancer, CD8^+^ T cells differentiate along an exhaustion trajectory, characterized by poor cytolytic function and expression of coinhibitory receptors, including PD-1^3^. “Stem-like”, “(pre)dysfunctional” or “progenitor exhausted (Tpex)” cells lie early in this trajectory, marked by co-expression of transcription factor TCF-1 and PD-1, next to CXCR5 and SLAMF6^4,5^. Stem-like CD8^+^ T cells are of particular interest, since they respond to PD-1-targeted immune checkpoint blockade (ICB) to generate functional CTLs in mouse models and are associated with ICB response in human cancer^6^.

Recent meta-analyses of mRNA- and TCR sequencing datasets have mapped stem-like CD8^+^ T cells at the branchpoint of differentiation trajectories towards either the functional CTL effector state or the terminally exhausted state^7,8^. Stem-like cells are formed during both acute and chronic infection, consistent with the concept that they can differentiate into either cell state^9,10^. To improve immunotherapy strategies, it is crucial to know how stem-like CD8^+^ T cells can be instructed to become competent CTL effectors^3,6^.

We here test the hypothesis that stem-like CD8^+^ T cells lie on a potentially productive CTL effector differentiation trajectory that can be completed when CD4^+^ T-cell help signals are provided^11^. During priming in lymph nodes (LNs), CD4^+^ T cells can deliver “help” for the CTL response, via conventional (c)DCs. CD4^+^ and CD8^+^ T cells are initially activated separately, in distinct regions of the LN by migratory cDCs. Subsequently, activated CD4^+^ and CD8^+^ T cells can recognize their respective antigens on the same cDC1. In this cellular scenario, help is relayed from CD4^+^ T cells to CD8^+^ T cells via cDC1s that though this “licensing” acquire optimal antigen cross-presentation ability and upregulate critical costimulatory ligands, cytokines and chemokines^12–14^. Helped CD8^+^ T cells undergo optimal CTL effector and effector memory differentiation which are pivotal for tumor control^15,16^. Conversely, “helpless” CTLs, i.e. CD8^+^ T cells primed in absence of CD4^+^ T-cell help, have typical hallmarks of CTL dysfunction^11,15,17^.

Inflammatory signals, in particular IFN-I, makes a key contribution to cDC1 licensing^18,19^. IFN-I acts in synergy with CD4^+^ T-cell mediated CD40 signaling to induce the gene expression program of the licensed cDC1^20^. Thus, innate signaling by IFN-I can partially compensate for CD4^+^ T-cell help, but it is not sufficient to allow for optimal CTL effector (memory) differentiation^13,21^. This is particularly relevant in cancer, where IFN-I signaling is often disabled^22^. In chronic LCMV infection, depletion of CD4^+^ T cells reduces formation of virus-controlling effector CTLs, but not stem-like cells^23,24^. Based on these concepts, we have postulated that stem-like cells as found in chronic infection and cancer may result from helpless priming^11^. Here, we tested the impact of CD4^+^ T-cell help on formation and fate of stem-like CD8^+^ T cells in a well-defined vaccination model, which was previously used to define how CD4^+^ T-cell help improves function of CD8^+^ T-cell effector and memory cells^15,16^. We demonstrate that PD-1^+^TCF-1^+^ stem-like CD8^+^ T cells lie on a potentially productive effector differentiation trajectory that is completed when help signals are delivered by vaccination. However, in a tumor context, help delivery and CTL effector differentiation are impeded, even when a CD4^+^ T-cell response to tumor-intrinsic epitopes takes place.

## METHODS

### Mice

Female 8-10-week-old C57BL/6JRj mice obtained from Janvier Laboratories and OT-I (Tg(TcraTcrb)1100Mjb) mice on a CD45.1^+^ C57BL/6 background (bred in-house) were housed in individually ventilated cages under specific pathogen free conditions. Animal experiments were approved by the Experimental Animal Committee of Leiden University Medical Center and performed in accordance with national and European regulations. Cages were allocated randomly to experimental groups.

### Vaccination

For intra-epidermal DNA vaccination, hair was removed from the right hind leg with depilating cream (Veet, Reckitt Benckiser) on day 0 and on days 0, 3, and 6, 15 μl of a 2 mg/ml plasmid DNA solution in H_2_O was applied into the hairless skin of anesthetized mice with a Permanent Make Up tattoo device (Cheyenne, MT Derm GmbH), using a sterile disposable nine-needle bar with a needle depth of 1 mm and oscillating at a frequency of 100 Hz for 45 s, as described^26,27^. Plasmid DNA vaccines used were HELP-E7SH (Help) and E7SH DNA (No Help)^25,26^, OVA-Help and OVA-No Help^15^ and Rpl18-Adpgk-Help vaccine. The latter was derived from the HELP-E7SH plasmid^26^ by replacing the E7SH sequence with codon-optimized sequences for mutated Adpgk (cacctggaactcgcgtccatgaccaacatggagctgatgtc-ctccatcgtgcatcag) and mutated Rpl18 (aaagcaggaggcaagattctgaccttcgacagactggcactggagagcccaaaa) epitopes^27^, separated by an alanine linker.

### Tumor challenge

The murine MC38-L colorectal carcinoma cell line was obtained from Leiden University Medical Center and has been characterized regarding neo-epitope expression^27–29^. The MC38-HELP cell line was created by transducing MC38-L cells with lentivirus encoding EF1a-HELP-3xFLAG-IRES-eGFP (**Supplementary Methods**). Mice were injected subcutaneously in the right flank with 300.000 live MC38 or MC38-HELP cells (LUMC) in 100 μl Hank’s buffered salt solution (HBSS, Gibco). Tumor sizes were measured 3 times per week by caliper, and a tumor volume of 1000 mm^3^ was maintained as humane endpoint.

### Tissue processing and flow cytometry

Peripheral blood cells were obtained by tail vein bleeding in Microvette CB300 LH tubes (Sarstedt) and erythrocytes were removed by incubating twice in Red Blood Cell lysis buffer (Santa Cruz) for 1 min at room temperature. Spleens were passed through a 70 μm cell strainer (BD Falcon) and erythrocytes were lysed for 5 min at room temperature. BM was obtained by gently crushing the bone of the right tibia with a mortar while adding Iscoves’ Modified Dulbecco Medium (IMDM, Lonza) + 10% Fetal Calf Serum (FCS) (Gibco). BM was passed through a 70 μm cell strainer, and erythrocytes were lysed for 5 min at room temperature. Right (dLN) and left (ndLN) inguinal lymph nodes were digested into single cell suspensions with 100 μg/ml Liberase TL (Roche) in Dulbecco’s Modified Eagle Medium (DMEM, Gibco) for 30 min at 37°C and passed through a 70 μm cell strainer. Tumors were isolated after perfusion of sacrificed mice with 15 ml phosphate buffered saline (PBS) with 2 mM ethylenediaminetetraacetic acid (EDTA, Santa Cruz). Tumors were mechanically disaggregated by scalpel and digested into single cell suspensions with 100 μg/ml Liberase TL (Roche) with 10 μg/ml DNAse (Sigma) in DMEM (Gibco) for 30 min at 37°C. Tumors were passed through a 70 μm cell strainer. For flow cytometry, cells were washed with FACS buffer (PBS + 2% FCS) and stained for 1 h at room temperature with 1:200 PE-conjugated I-Ab/AKFVAAWTLKAA (PADRE) tetramers (NIH) or for 30 min on ice with 1:1000 LIVE/DEAD Near IR dye (Invitrogen) or 1:500 Zombie UV (Biolegend), 1:100 APC-conjugated H-2D^b^/E7_49-57_ tetramers, APC-conjugated H-2K^b^/OVA_257-264_ tetramers or APC-conjugated H-2K^b^/KILTFDRL (mutated Rpl18) tetramers (produced in-house at LUMC) and monoclonal antibodies (mAbs) for surface staining (**Supplementary Methods**). For intracellular staining, cells were fixed and permeabilized with the FOXP3/transcription factor staining buffer set (eBioscience), according to the manufacturer’s protocol and stained with antibodies for 30 min on ice (**Supplementary Methods**). Flow cytometry was performed using a 3-laser or 5-laser Aurora (Cytek) and SpectroFlow software.

### *Ex vivo* restimulation

*Ex vivo* restimulation of splenocyte- or tumor cell suspensions was performed as described ^27^. In brief, cells of the DC line D1 were seeded in a 96-wells plate at 50.000 cells per well in conditioned medium and loaded with 1 mg/ml mutated Rpl18 and mutated Adpgk peptide or left unloaded, and incubated overnight at 37°C. The next day, 500.000 splenocytes or tumor cells were seeded on D1 cells and preincubated for 1 h at 37°C. Subsequently, GolgiPlug (BD Biosciences) was added at a final dilution of 1:1000, and left for 5 h at 37°C until analysis.

### Adoptive cell transfer

OT-I CD8^+^ T cells, carrying a transgenic Vα2/Vβ5 TCR recognizing OVA_257-264_/H-2K^b^, were isolated to more than 90% purity from spleens of naïve OT-I;CD45.1^+^ C57BL/6 mice using a negative selection MACS CD8α^+^ T-cell Isolation Kit (Miltenyi). OT-I cells were retro-orbitally (r.o.) injected at 10^5^ cells per mouse into WT CD45.2^+^ C57BL/6 mice on day -1, after which mice were vaccinated with OVA-No Help construct^15^ on day 0, 3 and 6. On day 10, CD45.1^+^ CD44^+^ SLAMF6^+^ helpless CD8^+^ T cells were sorted from dLN on a 3-laser FACSAria II (BD Biosciences). Cells were r.o. injected at 10^4^ cells per mouse into WT C57BL/6 mice, which were vaccinated once with OVA-Help or OVA-No Help on the next day.

### Flow cytometry data analysis

Flow cytometry data were analyzed with OMIQ software and FlowAI^30^ was used to remove events showing anomalies in flow rate, signal acquisition and dynamic range. After manual gating of tetramer^+^ CD8^+^ T cells (**Supplementary Figure 1A**) and conventional marker expression analysis, subsampling was performed to select maximally 150 tetramer^+^ CD8^+^ T cells from each sample. Samples containing <5 total tetramer^+^ CD8^+^ T cells were excluded from further analysis. FlowSom^31^ was used for K-means clustering of tetramer^+^ CD8^+^ T cells, including all markers except live/dead, CD4, CD8, FOXP3 and tetramer. Optimal t-distributed stochastic neighbor embedding (opt-SNE)^32^ was used for dimension reduction and visualization of the data, using the same markers. Wanderlust^33^ was used for trajectory analysis, using the same markers except Ki67, and using Cluster 10 (CD44^-^CD62L^+^ naive cells) as starting point. FlowSom clustering of all tetramer^+^ CD8^+^ cells from dLN at day 2-10 and 3 naïve LNs was done for the analysis of antigen-specific CD8^+^ T-cell populations on consecutive days after vaccination. Responding tetramer^+^ CD8^+^ T cells were selected for analysis by including clusters that showed <20% frequency in naïve LNs and a change in frequency between different time points after vaccination (**Supplementary Figure 3D**).

### Statistical analysis

Data was analyzed with GraphPad Prism software using two-way ANOVA and Sidak’s multiple comparisons test, unless indicated otherwise. Error bars in figures indicate SD. A P value < 0.05 was considered statistically significant; *P < 0.05, **P < 0.005, ***P < 0.001 and ****P < 0.0001. Sample size was calculated a priori using G*Power software, using previous results of E7-specific CD8^+^ T-cell frequencies between Help and No Help^15^ for power calculations.

### Sample and library preparation for scRNAseq and TCRseq

Mice were vaccinated under Help or No Help conditions at day 0, 3 and 6 and were sacrificed at day 5 or day 10 after first vaccination, after which dLN and ndLN were isolated (**Supplementary Figure 5**). For each experimental condition, LNs from 3 individual mice were pooled to reduce biological variability, resulting in 8 samples (**Supplementary Figure 6A**). Samples were stained with the following mAbs: 1:100 CD19-PerCP/Vio700 (clone REA749, Miltenyi), 1:100 CD3-BV421 (clone 17A2, Biolegend), 1:100 CD44-PE/Cy7 (clone IM7, eBioscience), 1:100 CD62L-FITC (clone MEL-14, eBioscience). For each sample, activated T cells (CD19^-^CD3^+^CD44^+^CD62L^-^) were sorted on a 3-laser FACSAria II (BD Biosciences) and labeled with a distinct hashtag oligonucleotide (HTO)-conjugated mAb to MHC-I/CD45 (TotalSeq C0301 to C0307 and C0309, Biolegend), after which all 8 samples were pooled. scRNAseq libraries were prepared using a 10X Chromium single cell 5’ v2 chemistry kit (10X Genomics) andl TCRseq libraries were prepared using a 10X Chromium single cell V(D)J enrichment kit for mouse T cells (10X Genomics), according to the manufacturer’s protocols. Sequencing was performed on a NovaSeq6000 system (Illumina). A total of 25,000 cells was used for sequencing, with a median sequencing depth of 11,196 reads per cell.

### scRNAseq analysis

Analysis of scRNAseq and TCRseq data was performed using CellRanger (10X Genomics, v6.0.1) and Seurat (v4.3.0) in R (v4.2.3). For details, see **Supplementary Methods**. In summary, scRNAseq and TCRseq reads were aligned to the mouse reference genome mm10, TCR and BCR genes were removed from the gene expression count matrix, and the dataset was analyzed using the standard Seurat pipeline. HTO demultiplexing was performed (**Supplementary Figure 6B**) and dataset was filtered based on number of RNA features, number of RNA counts, mitochondrial gene expression and expression of genes associated with B cells or APCs. Cell cycle scores for G2/M and S phase were assigned based on expression of G2/M and S phase genes, and these scores were regressed out during clustering. CD8^+^ T-cell clusters were selected on presence of *Cd8a, Cd8b1* and absence *Cd4* transcripts (**Supplementary Figure 6C**). Within the CD8^+^ T-cell subset, number of cells per HTO was determined (**Supplementary Figure 6D**). HTOs containing <30 CD8^+^ T cells were excluded from further analysis. Cluster 7, consisting of 52 CD8^+^ T cells that showed only cell cycle genes as markers, was excluded, since cell cycle genes were regressed out during clustering. TCRseq analysis was performed using Loupe VDJ Browser (v5.0.0, 10X Genomics). Expanded clones were identified as TCR clonotypes containing 2 or more cells. TCR sequences from Loupe VDJ Browser, and UMAP projection and cluster annotation from Seurat were visualized together using Loupe Browser (v6.4.1, 10X Genomics). To quantify expanded clonotypes per differentiation cluster, only cells that grouped together in the UMAP with their assigned cluster were included. Seurat and Monocle3 (v1.3.1) in R were used for trajectory analysis. Ingenuity Pathway Analysis (Qiagen) was used for functional classification of differentially expressed genes between Cluster 2 and Cluster 3. For comparison of our scRNAseq data to published gene signatures, top 200 marker genes of TCF-1^+^ progenitor (cluster 9) and effector (cluster 6) CD8^+^ T cells from chronic and acute LCMV infection were downloaded from Table S5 from Pritykin *et al*. 2021^7^. These top 200 genes were used to calculate a module score for each cell in our dataset. The same method was performed using top 100 marker genes of stem-like CD8^+^ T cells from the TdLN, (cluster 4) as published by Connolly *et al.* 2021^34^, which were kindly provided to us by the authors.

## RESULTS

### Stem-like CD8^+^ T cells predominate after priming in absence of CD4^+^ T-cell help

To compare CD8^+^ T-cell responses in presence or absence of CD4^+^ T-cell help, we use two DNA vaccines: one that encodes the immunodominant E7_48-57_ peptide of human papilloma virus (HPV) that only primes E7-specific CD8^+^ T cells (No Help) and one that additionally encodes HPV-unrelated MHC-II restricted helper epitopes P30/PADRE and also primes CD4^+^ T cells^25,26^ (**Figure 1A**). The plasmid DNA is “tattooed”’ into the epidermis, resulting in antigen expression in keratinocytes and efficient T-cell priming by migratory cDCs in the skin-draining inguinal LN^25,35^. Mice are vaccinated at day 0, 3 and 6, after which the antigen-specific CD8^+^ T-cell response is followed by H-2D^b^/E7_48-57_ (E7) tetramer staining (**Supplementary Figure 1A-C**). This is a transparent model to investigate the impact of CD4^+^ T-cell help during priming on long-term CD8^+^ T-cell fate, since the antigen is only transiently present during priming^35,36^ and the impact of conventional antigen-specific CD4^+^ T cells is tested by their vaccine-based engagement in the response^25,26^.

**Figure 1.**
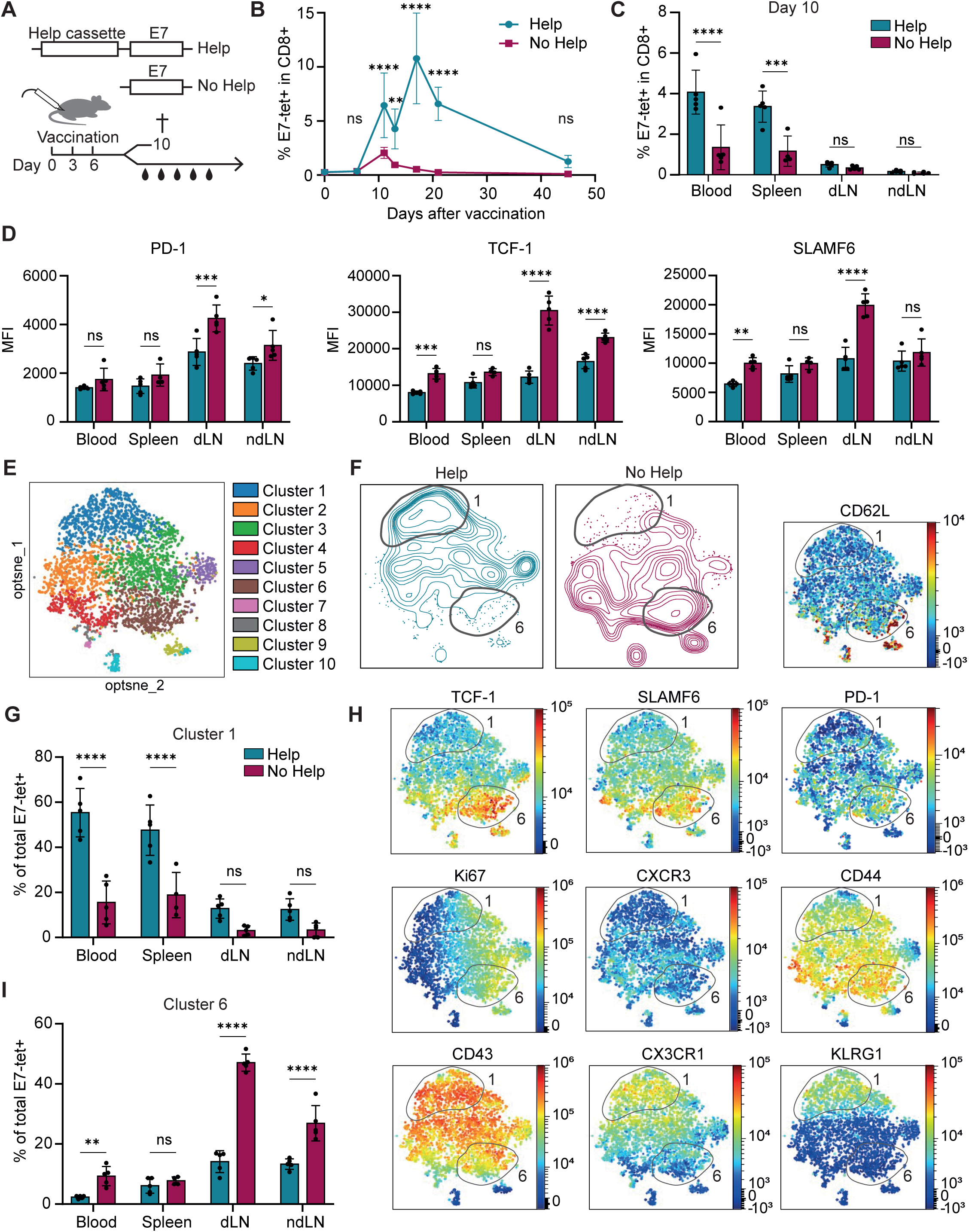
Stem-like CD8^+^ T cells predominate after priming in absence of CD4^+^ T-cell help. **(A)** Vaccination model. Plasmid DNA, encoding non-functional HPV E7 protein (E7) and helper epitopes (P30, PADRE) (Help) or not (No Help) is tattooed into the epidermis of mice at day 0, 3 and 6 to raise an H-2D^b^/E7_48-57_ (E7)-specific CD8^+^ T-cell response. (**B-I**) Mice were vaccinated under Help or No Help conditions and the response of E7-tetramer^+^ CD8^+^ T cells was analyzed by flow cytometry with antibodies to differentiation markers. (**B-C**) Percentage of E7-tetramer^+^ CD8^+^ T cells in blood at the indicated days after the first vaccination (B) and in indicated organs at day 10 (C). (**D**) Median Fluorescence Intensity (MFI) of indicated markers on E7-tetramer^+^ CD8^+^ T cells at day 10. (**E, F**) Opt-SNE visualization of E7-tetramer^+^ CD8^+^ T cells at day 10. FlowSom clustering of cells from all experimental settings (E) and distribution of cells from the two vaccination settings (F). Circled areas indicate clusters 1 and 6. (**G**) Frequency of E7-tetramer^+^ CD8^+^ T cells in Cluster 1 per organ after Help versus No Help vaccination. (**H**) Expression of differentiation markers on cumulative E7-tetramer^+^ CD8^+^ T cells from all experimental settings visualized by opt-SNE. (**I**). Frequency of E7-tetramer^+^ CD8^+^ T cells in Cluster 6 per organ after Help versus No Help vaccination. Data are from 1 experiment, representative of at least 2 independent experiments with n=5 mice per group. Error bars indicate SD. Statistical significance was calculated by two-way ANOVA and Sidak’s multiple comparisons test. *P < 0.05, **P <0.005, ***P < 0.001 and ****P < 0.0001. See also Supplementary Figure 1.

As previously shown^15,25^, the E7-specific CD8^+^ T-cell response to No Help vaccination was lower and of shorter duration than the response to Help vaccination (**Figure 1B**). At day 10, the frequency of E7-specific CD8^+^ T cells was significantly lower in the No Help setting than in the Help setting in blood and spleen (**Figure 1C**). Moreover, in the No Help setting PD-1, TCF-1 and SLAMF6 that identify the progenitor exhausted/stem-like state^6,24,37^ were expressed at higher levels on E7-specific CD8^+^ T cells in the dLN, which is the priming site (**Figure 1D**).

Next, we used an extensive antibody panel to distinguish E7-specific CD8^+^ T-cell differentiation states^2,38^ after Help or No Help vaccination in blood, spleen, dLN and ndLN. At day 10 after vaccination, clustering analysis and dimension reduction on cumulative E7-tetramer^+^ CD8^+^ T cells resulted in 10 clusters (**Figure 1E**), which were differentially distributed between vaccination settings (**Figure 1F, Supplementary Figure 1D**) and between organs (**Supplementary Figure 1D, E**). Cluster 1 cells were higher in frequency among tetramer^+^ CD8^+^ T cells (**Figure 1G**) and total CD8^+^ T cells (**Supplementary** Figure 1F) after Help than after No Help vaccination and were mostly found in the circulation, i.e. in blood and spleen. Cluster 1 cells were characterized by low expression of CD62L and TCF-1 and high expression of KLRG1 and CX3CR1 (**Figure 1H**), consistent with a terminally differentiated, short-lived effector CTL state^39,40^. In lymph nodes, Cluster 6 cells were higher in frequency among tetramer^+^ CD8^+^ T cells (**Figure 1I**) and total CD8^+^ T cells (**Supplementary Figure 1F**) in the No Help setting than in the Help setting. Cluster 6 cells were characterized by high expression of TCF-1, SLAMF6 and PD-1 and low expression of KLRG1 and CX3CR1 (**Figure 1H**), indicating the stem-like phenotype^6,24,37^. Consistent with vaccination rather than chronic antigen exposure, we did not observe exhausted CD8^+^ T cells with a PD-1^high^TIM3^+^TCF-1^-^ phenotype^4^ (**Supplementary Figure 1G**). Together, these results indicate that in presence of CD4^+^ T-cell help during priming, CD8^+^ T cells can terminally differentiate into CTL effector cells that distribute systemically through blood and spleen. In absence of CD4^+^ T-cell help, primed CD8^+^ T cells accumulate in the PD-1^+^TCF-1^+^ stem-like state in LNs.

### CD4^+^ T-cell help shifts the CD8^+^ T-cell differentiation spectrum towards the effector state

To analyze the effects of CD4^+^ T-cell help on the CTL differentiation spectrum, we mapped the antigen-specific CD8^+^ T-cell clusters as defined in Figure 1 using the Wanderlust algorithm that aligns cells to a unified trajectory based on overlap in marker expression^33^. This procedure generated a CTL differentiation trajectory ranging from the naïve state (CD44^-^CD62L^+^) to the short-lived effector state (KLRG1^+^CX3CR1^+^) (**Figure 2A**). After Help vaccination, CD8^+^ T cells in dLN, spleen and blood had a higher mean trajectory score than after No Help vaccination (**Figure 2B**), indicating that on average, CD4^+^ T-cell help shifts the differentiation state of CD8^+^ T cells towards completion of the trajectory.

**Figure 2.**
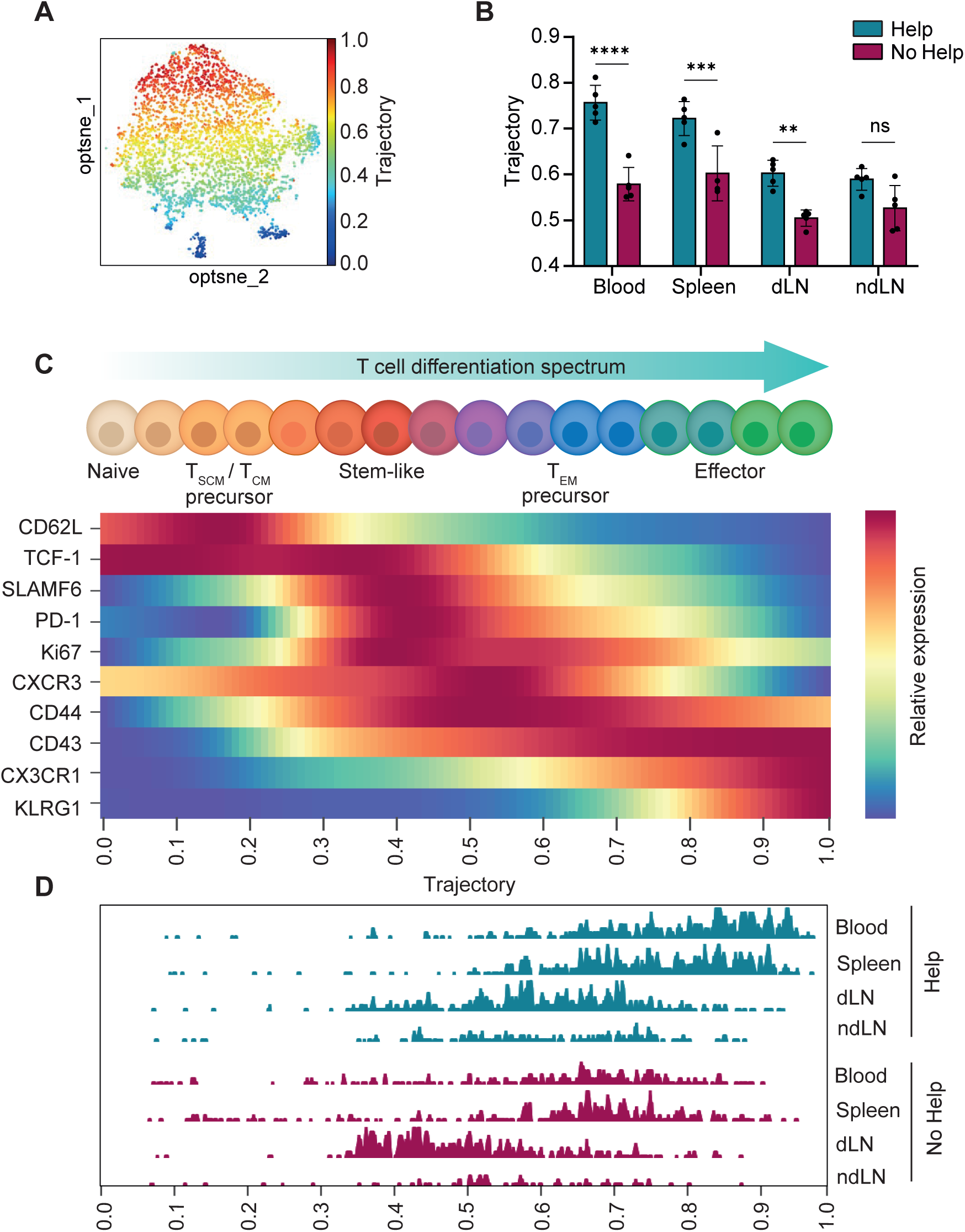
CD4^+^ T-cell help shifts the CD8^+^ T-cell differentiation spectrum towards the effector state. Mice were vaccinated under Help or No Help conditions and analyzed at day 10 as outlined in Figure 1 (same experiment). (**A**) Wanderlust trajectory shown as color gradient within opt-SNE visualization of all E7-tetramer^+^ CD8^+^ T cells from all organs. (**B**) Mean trajectory score of E7-tetramer^+^ CD8^+^ T cells from indicated organs. (**C**) CD8^+^ T-cell differentiation states as defined in literature were mapped to the trajectory axis based on marker expression. Heat map showing relative expression levels of differentiation markers on E7-tetramer^+^ CD8^+^ T cells. For each marker, relative expression was determined based on lowest and highest expression within the E7-tetramer^+^ CD8^+^ T-cell population. (**D**) Histograms showing distribution of E7-tetramer^+^ CD8^+^ T cells from indicated organs after Help or No Help vaccination along the differentiation trajectory. Data are from 1 experiment, representative of 2 independent experiments with n=5 mice per group. Error bars indicate SD. Statistical significance was calculated by two-way ANOVA and Sidak’s multiple comparisons test. *P < 0.05, **P< 0.005, ***P < 0.001 and ****P <0.0001. See also Supplementary Figure 2.

To visualize the calculated trajectory along the progressive differentiation path as described in literature^1,2,38^, we mapped protein expression of differentiation markers of naïve (CD44^-^CD62L^+^TCF-1^+^), T stem cell memory (T_SCM_)/T_CM_ precursor (CD44^+^CD62L^+^TCF-1^+^), stem-like (PD-1^+^TCF-1^+^SLAMF6^+^), T_EM_ precursor (CD44^+^CD62L^-^) and effector (KLRG1^+^CX3CR1^+^) cell states against the trajectory (**Figure 2C, Supplementary Figure 2**). PD-1 and SLAMF6 were upregulated halfway through the trajectory, in CD8^+^ T cells that retained TCF-1 expression, corroborating that PD-1^+^TCF-1^+^SLAMF6^+^ stem-like cells are in an early differentiation state. These cells highly expressed proliferation marker Ki67, in agreement with literature data^37,41,42^. Further differentiation along the trajectory towards effector cells was associated with downregulation of PD-1, TCF-1 and SLAMF6 (**Figure 2C**).

Next, we plotted antigen-specific CD8^+^ T cells, harvested from blood, spleen and LNs after Help or No Help vaccination, against this differentiation spectrum based on their trajectory score (**Figure 2D**). Blood and spleen harbored cells with a high trajectory score of 0.8-1.0 (**Figure 2D**), corresponding to a KLRG1^+^CX3CR1^+^ phenotype (**Figure 2C**) after Help vaccination, while this population was all but lacking after No Help vaccination. In LNs, cells had a lower trajectory score ranging from 0.3 to 0.8 (**Figure 2B, 2D**), predominated by the PD-1^+^TCF-1^+^SLAMF6^+^ phenotype (**Figure 2C**), both after Help and No Help vaccination. However, after Help vaccination the LN cells had differentiated further than after No Help vaccination (**Figure 2B, 2D**).

We conclude that antigen-primed CD8^+^ T cells can differentiate into CD44^+^CD62L^+^ T_CM_ precursor and PD-1^+^TCF-1^+^SLAMF6^+^ stem-like populations irrespective of CD4^+^ T-cell help, but require CD4^+^ T-cell help to become CD44^+^CD62L^-^ T_EM_ precursor and KLRG1^+^CX3CR1^+^ short-lived CTL effector cells.

### Stem-like CD8^+^ T cells turn into effector cells upon receiving CD4^+^ T-cell help

If stem-like CD8^+^ T-cells are indeed incompletely differentiated, one would expect them to arise earlier after priming than help-dependent effector CTLs. To test this assumption, we vaccinated mice under Help or No Help conditions and analyzed antigen-specific CD8^+^ T-cells in the dLN at multiple successive days after the first vaccination (**Figure 3A**). After Help vaccination, E7-specific CD8^+^ T cells strongly increased in number from day 6 onwards, while after No Help vaccination the response was lower and more belated (**Figure 3B**). To exclude background tetramer staining of naïve CD8^+^ T cells, especially at early time points, we only included clusters that increased in frequency after vaccination (**Supplementary Figure 3**). Clustering and dimension reduction of cumulative responding CD8^+^ T cells indicated distinct phenotypes in Help and No Help settings, as expected (**Figure 3C**). We dissected responder CD8^+^ T cells into stem-like (TCF-1^+^SLAMF6^+^) and effector (GZMB^+^) subsets (**Figure 3D, E**) and analyzed the generation of these two populations over time. After Help vaccination, the stem-like population initially predominated, but from day 6 onwards it decreased, while the effector population increased (**Figure 3F**). After No Help vaccination, in contrast, the stem-like population steadily increased to form the majority of the E7-specific CD8^+^ T-cell pool in the dLN, while the effector cell population remained small (**Figure 3G**). These results indicate that stem-like CD8^+^ T cells are generated early after priming, in both presence and absence of CD4^+^ T-cell help. However, only in presence of CD4^+^ T-cell help, these cells in time differentiate further into effector CTLs.

**Figure 3.**
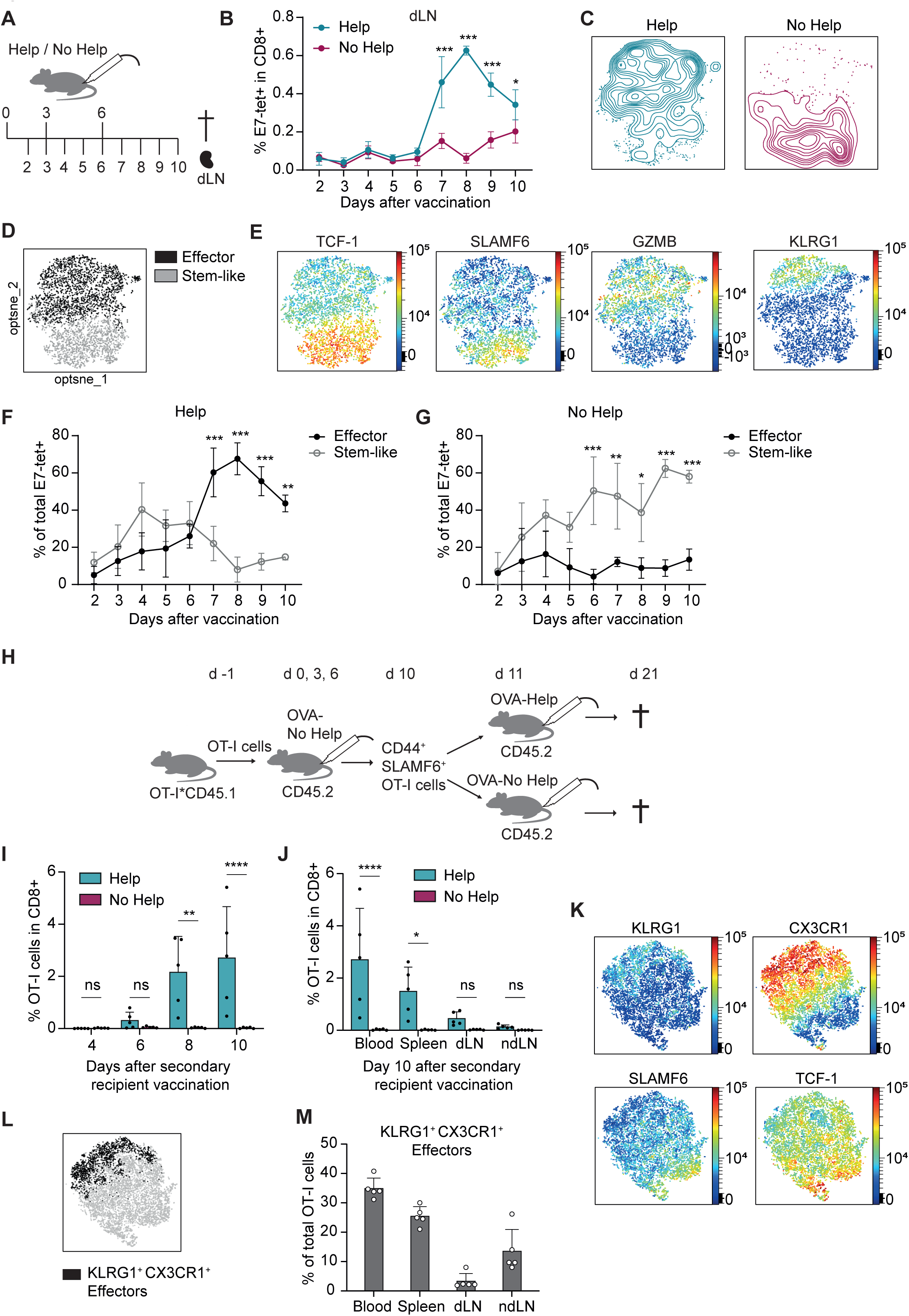
Formation of effector but not stem-like CD8^+^ T cells relies on CD4^+^ T-cell help. **(A)** Experimental setup for panels B-G. Mice received Help or No Help vaccination at day 0, 3 and 6, and different cohorts were sacrificed at day 2-10 after the first vaccination, after which E7-specific CD8^+^ T cells in the dLN were analyzed by flow cytometry with the differentiation marker panel. (**B**) Percentage of E7-tetramer^+^ CD8^+^ T cells in dLN after Help or No Help vaccination. (**C-E**) Opt-SNE visualization of all responding E7-tetramer^+^ CD8^+^ T cells (see Supplementary Figure 3) in the dLN at all time points after Help or No Help vaccination. (**C**) Distribution of responder CD8^+^ T cells from Help versus No Help vaccination settings. (**D**) Visualization of stem-like (TCF-1^+^SLAMF6^+^) and effector (GZMB^+^) clusters. (**E**) Visualization of differentiation marker expression. (**F, G**) Frequency of stem-like and effector populations as defined in C-E within total responder E7-tetramer^+^ CD8^+^ T cells at different time points after vaccination. (**H**) Experimental setup for panels I-M. CD45.2^+^ naïve mice received 10^5^ CD45.1^+^ OT-I T cells and OVA-No Help vaccine. At day 10, CD45.1^+^ OT-I T cells of the were flow cytometrically sorted from dLNs on CD44^+^SLAMF6^+^ phenotype (Supplementary Figure 4) and transferred at 10^4^ cells per mouse to CD45.2^+^ secondary recipients that received OVA-Help or OVA-No Help vaccine on the next day. (**I, J**) Percentage of OT-I (CD45.1^+^OVA-tetramer^+^ CD8^+^) T cells in secondary recipients after Help or No Help vaccination, as measured in blood at different time points (I) and in indicated organs at day 10 (J). (**K, L**) Opt-SNE visualization of OT-I T cells in dLN of secondary recipients at day 10 after Help vaccination. (**K**) Expression of differentiation markers across opt-SNE. (**L**) Visualization of the KLRG1^+^CX3CR1^+^ effector population. (**M**) Frequency of KLRG1^+^CX3CR1^+^ effector populations within the OT-I T cell population in indicated organs. Data from B-G and I-M are each from 1 experiment, representative of 2 independent experiments with n=3-5 mice per group. Error bars indicate SD. Statistical significance was calculated by two-way ANOVA and Sidak’s multiple comparisons test. *P < 0.05, **P < 0.005, ***P < 0.001 and ****P < 0.0001. See also Supplementary Figure 3 and 4.

It is an important question whether stem-like CD8^+^ T cells primed in absence of CD4^+^ T-cell help can still complete their effector differentiation when help signals are delivered after priming. To test this, we used adoptive transfer with OT-I T cells that express a transgenic TCR recognizing the ovalbumin (OVA)_257/264_/H-2K^b^ complex (**Figure 3H**). CD45.2^+^ mice received naïve CD45.1^+^ OT-I cells and OVA-No Help vaccine encoding the OVA_257/264_ epitope. At day 10, CD45.1^+^CD44^+^SLAMF6^+^ stem-like OT-I cells were flow sorted from pooled dLNs (**Supplementary Figure 4A**) and transferred into CD45.2^+^ secondary recipient mice. The next day, these mice received either control OVA-No Help or test OVA-Help vaccine with the same helper epitopes as the E7 vaccine. Transferred stem-like OT-I T cells clonally expanded upon OVA-Help vaccination, but not upon OVA-No Help vaccination (**Figure 3I, J, Supplementary Figure 4B**). After Help vaccination, a proportion of the transferred OT-I T cells displayed the help-dependent KLRG1^+^CX3CR1^+^ effector phenotype (**Figure 3K, L**), primarily in blood and spleen (**Figure 3M**). Together, these results demonstrate that helpless stem-like CD8^+^ T cells can still respond to help signals after initial priming by both expansion, CTL effector differentiation and systemic dissemination.

### scRNAseq and TCRseq support that stem-like CD8^+^ T cells become CTL effectors upon CD4^+^ T-cell help delivery

To test directly whether addition of CD4^+^ T-cell help signals leads to the transition of stem-like CD8^+^ T cells into effector CTLs, we performed single cell RNA sequencing (scRNAseq) and TCR sequencing (TCRseq). For this purpose, mice received E7-Help or E7-No Help vaccine and at day 5 or day 10, activated CD3^+^CD44^+^CD62L^-^ T cells were sorted by flow cytometry from both dLN and ndLN (**Supplementary Figure 5**). This resulted in eight samples that were labeled with distinct hashtag oligonucleotides (HTO)-conjugated antibodies before performing scRNAseq and TCRseq on pooled cells (**Supplementary Figure 6A, B**). For analysis, we used CD8^+^ T-cells isolated from dLNs at days 5 and 10 after Help vaccination and at day 10 after No Help vaccination, since these HTO groups yielded sufficient CD8^+^ T-cell numbers as identified based on *Cd8a* and *Cd8b1* expression (**Supplementary Figure 6C, D**).

Whole transcriptome-based clustering of 3051 CD8^+^ T cells pooled from these three experimental conditions identified seven distinct cell states (**Figure 4A**), based on specific transcripts (**Supplementary Table 1**). Naive cells (Cluster 0) and T_CM_ cells (Cluster 1) were identified by co-expression of *Sell* (encoding CD62L), *Il7r* and *Ly6c2*, while naive cells were expressed higher levels of *Lef1* and *Ccr7* and T_CM_ cells expressed *Eomes* (**Figure 4B**). Stem-like cells (Cluster 2) were identified by high expression of *Tcf7* (encoding TCF-1), *Id3* and *Slamf6,* next to *Sell*, *Lef1* and *Ccr7*. CTL effector cells (Cluster 3) were characterized by expression of *Gzmb*, *Gzmk*, *Id2* and *Tbx21* (encoding T-bet) and also expressed various killer lectin receptor (KLR) genes such as *Klrc1* and *Klrk1* and proliferation-associated genes including *Mki67, Tk1 and Kif11*. Some smaller clusters were characterized by expression of KLRs (Cluster 4), protein/RNA biosynthesis genes (Cluster 5), and IFN-I responsive genes (Cluster 6).

**Figure 4.**
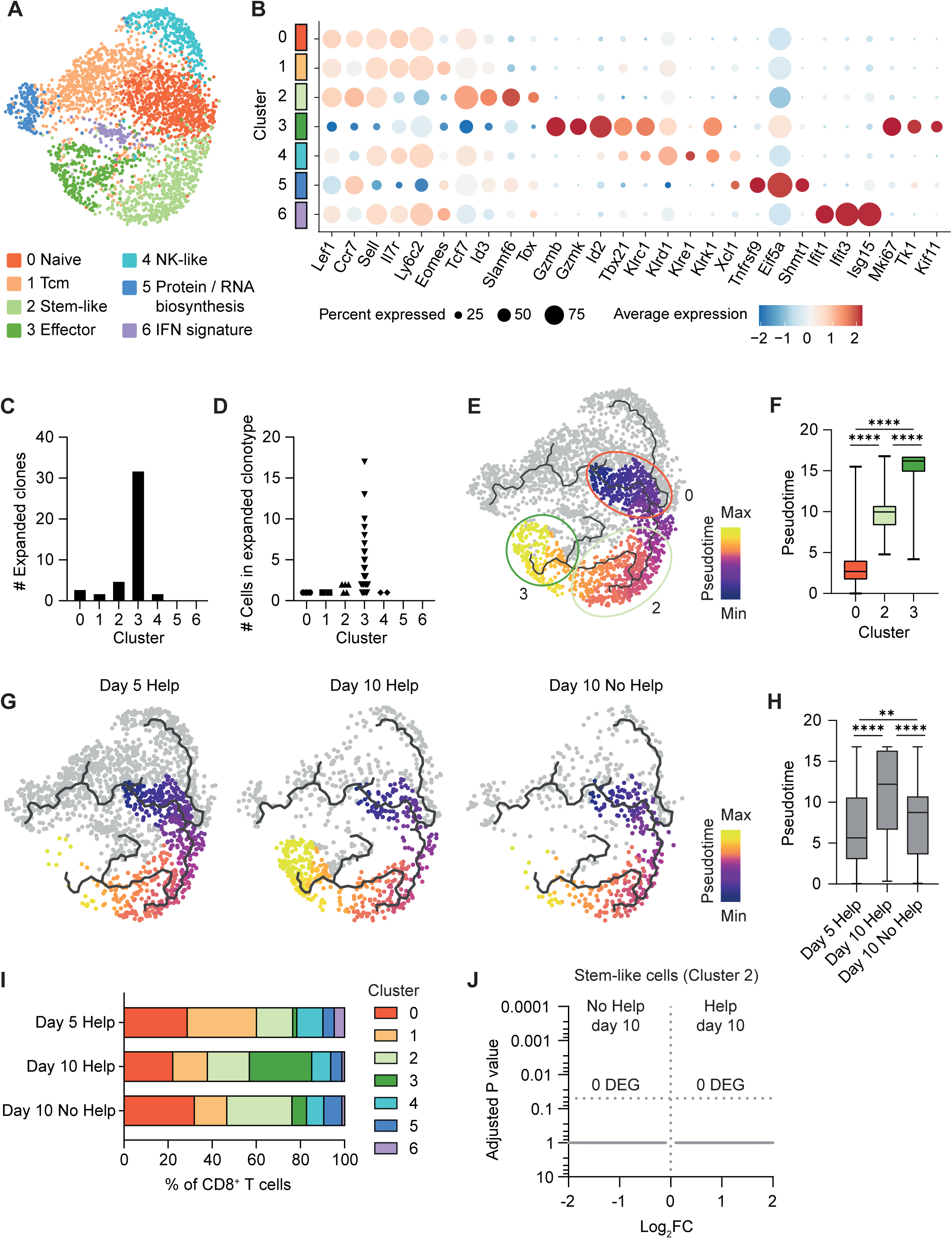
scRNAseq and TCRseq support conversion of stem-like CD8^+^ T cells into CTL effectors upon CD4^+^ T-cell help delivery. Mice (n=3 per group) were vaccinated under Help or No Help conditions and activated CD3^+^CD44^+^CD62L^-^ T cells were isolated by flow cytometry from dLN and ndLN at day 5 and day 10 (See Supplementary Figure 5). They were labeled with different HTOs per experimental group and subjected to 10x Genomics scRNAseq and coupled TCRseq (See Supplementary Figure 6). (**A**) UMAP projection and clustering of CD8^+^ T cells (defined in Supplementary Figure 6) from day 5 and day 10 Help and day 10 No Help vaccination conditions. (**B**) Dot plot showing level and frequency of CD8^+^ T-cell differentiation marker gene transcripts identified by scRNAseq per CD8^+^ T-cell cluster as defined in panel A. (**C, D**) TCR clonotype analysis. (**C**) Number (#) of distinct clonally expanded activated CD8^+^ T cells as deduced from TCRseq per cluster, based on ≥ 2 cells with the same TCR. (**D**) Number (#) of cells per expanded TCR clonotype per cluster (See also Supplementary Figure 7). (**E**) UMAP showing pseudotime within the differentiation trajectory branch from naïve to effector gene expression profile, as based on scRNAseq data from the cumulative day 5 Help and day 10 Help and No Help conditions. (**F**) Box plot showing pseudotime per cluster. Statistical significance was calculated by two-way ANOVA and Sidak’s multiple comparisons test. ****P < 0.0001. (**G**) UMAP showing pseudotime within the differentiation trajectory branch from naïve to effector gene expression profile, plotted separately per indicated experimental condition. (**H**) Distribution of CD8^+^ T cells over the different clusters (defined in A) per experimental condition. (**I**) Boxplot showing pseudotime per experimental condition. Statistical significance was calculated by two-way ANOVA and Sidak’s multiple comparisons test. **P < 0.005 and ****P <0.0001 (**J**) Volcano plot showing 0 differentially expressed genes (DEG) between CD8^+^ cells within Cluster 2 from the day 10 Help versus No Help settings.

Clonal expansion within each CD8^+^ T-cell cluster was assessed by TCRseq analysis. Expanded clones, defined as TCR clonotypes containing ≥ 2 cells, were quantified for each cluster (**Supplementary Figure 7A**). This analysis revealed that, in agreement with proliferation marker expression, the effector population (Cluster 3) displayed much higher clonal expansion than any of the other clusters, both by number of distinct expanded clones (**Figure 4C**, **Supplementary Figure 7B, C**) and number of individual cells per clonotype (**Figure 4D**). The great majority of expanded TCR clonotypes were exclusively present in Cluster 3, while some were shared between Clusters 2 and 3 (**Supplementary Figure 7C**).

Pseudotime analysis based on calculating gene expression changes associated with dynamic biological processes^43^ identified the stem-like population (Cluster 2) as an intermediate state between the naive (Cluster 0) and effector (Cluster 3) CD8^+^ T-cell differentiation states (**Figure 4E, F**). Importantly, effector CD8^+^ T cells lying at the end of the differentiation trajectory (Cluster 3) (**Figure 4E**) were abundantly present in the day 10 Help condition, but mostly lacking in the day 10 No Help condition (**Figure 4G**). Under Help conditions, Cluster 3 effector cells increased in frequency from day 5 tot day 10 after vaccination (**Figure 4G**). Accordingly, CD8^+^ T cells from the day 10 Help condition showed a higher median pseudotime score as compared to cells from the other two experimental conditions (**Figure 4H**). They also showed a higher number of distinct expanded clones (**Supplementary Figure 7D, E**), and a higher number of cells per TCR clonotype (**Supplementary Figure 7F**), in agreement with clonal expansion being most prominent in Cluster 3 (**Figure 4C, D**). Stem-like CD8^+^ T cells (Cluster 2) were generated both under No Help and Help conditions (**Figure 4G, I**), but predominated under No Help conditions at day 10 (**Figure 4I**). Importantly, Cluster 2 cells from Help and No Help conditions had indiscernible transcriptomes at day 10 (**Figure 4J**). Differential gene expression analysis between Cluster 2 and 3 followed by Ingenuity Pathway Analysis (IPA) showed that functional qualities of helped CD8^+^ T cells, such as cytotoxicity, cell migration and proliferation, were acquired upon differentiation from the stem-like to the effector state (**Supplementary Figure 8A-D, Supplementary Table 2**), in agreement with our earlier data^15^.

Taken together, these results show at the single cell level that the differentiation spectrum of CD8^+^ T cells primed in presence of CD4^+^ T-cell help includes both stem-like and effector cells, with effector cells expanding and predominating in time. In absence of help, the same stem-like CD8^+^ T cells are formed that do not further differentiate into effector cells.

### Stem-like CD8^+^ T cells that predominate after helpless priming transcriptionally resemble stem-like CD8^+^ T cells in chronic infection and cancer

Next, we determined how the stem-like and helped effector CD8^+^ T-cell states from our vaccination system relate to CD8^+^ T-cell states found in infection and cancer. We used primary data from mouse infection models generated by Pritykin *et al*. (**Figure 5A**), who showed that the TCF-1^+^ progenitor/stem-like state of CD8^+^ T cells is epigenetically and transcriptionally comparable in classical models of chronic infection and cancer in mice. This study^7^ and others^8,10^ also demonstrated that the stem-like CD8^+^ T-cell state lies at the basis of trajectories towards either effector CD8^+^ T cells, as found in acute infection, or terminally exhausted CD8^+^ T cells, as found in chronic infection and cancer (**Figure 5B**). We used the top 200 signature genes of the TCF-1^+^ progenitor state defined by Pritykin *et al.* (**Supplementary Table 3**) to determine similarity to Cluster 2 and Cluster 3 cells from our vaccination model by module scoring. The TCF-1^+^ progenitor signature was clearly enriched in the stem-like Cluster 2 cells predominating after No Help vaccination (**Figure 5C**). Furthermore, the gene signature of effector CTLs responding to acute LCMV infection (**Supplementary Table 3**) was enriched in effector Cluster 3 cells that predominate after Help vaccination (**Figure 5C**).

**Figure 5.**
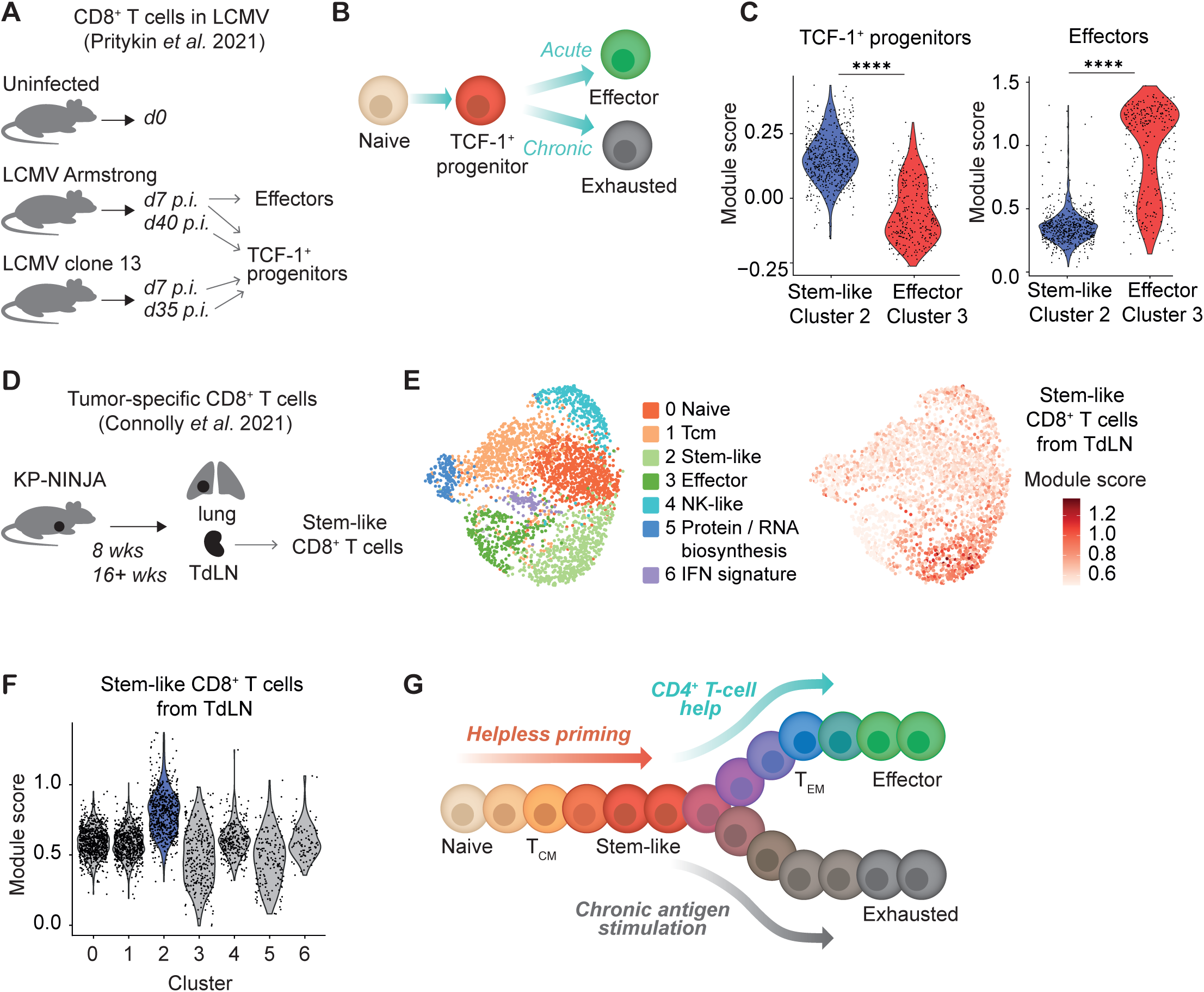
Stem-like CD8^+^ T cells that predominate after helpless priming transcriptionally resemble stem-like CD8^+^ T cells in chronic infection and cancer. **(A)** Outline of experimental conditions that generated the scRNAseq dataset from Pritykin *et al.*^8^, containing CD8^+^ T cells from uninfected mice and mice infected with either acute LCMV Armstrong or chronic LCMV clone 13 virus. TCF-1^+^ progenitors were found in all infected mice, while effector cells were only found at day 7 after LCMV Armstrong infection. (**B**) Differentiation model as proposed by Pritykin *et al*.^8^ and others^9,11^, where TCF-1^+^ progenitor cells lie at the branchpoint between effector and exhaustion differentiation trajectories, as followed by CD8^+^ T cells under conditions of acute or chronic antigen exposure. (**C**) Violin plots showing module scores of the top 200 marker genes for CD8^+^ TCF-1^+^ progenitor cells (cluster 9, left panel) and effector cells (cluster 6, right panel) from Pritykin *et al.*^8^ per CD8^+^ T cell in the stem-like Cluster 2 and helped effector Cluster 3 as defined in our vaccination model. Statistical significance was calculated by Mann-Whitney test. ****P < 0.0001. (**D**) Outline of experimental conditions that generated the scRNAseq dataset of Connolly *et al.*^37^. Tumor antigen-specific CD8^+^ T cells were isolated from tumors and TdLNs at an early (8 weeks) and late stage (16+ weeks) after induction of lung tumors the genetic KP-NINJA model. (**E, F**) Module score of the top 100 marker genes for stem-like CD8^+^ T cells from TdLN (cluster 4, Connolly *et al*.^37^) per CD8^+^ T cell present in Cluster 2 and 3 in our vaccination model (Figure 4), as shown by UMAP (E) and violin plots (F). (**G**) Proposed model for CD8^+^ T cell differentiation, wherein the PD-1^+^TCF-1^+^ stem-like state lies at the branchpoint of CD4^+^ T-cell help-dependent effector CTL differentiation and chronic antigen-driven CD8^+^ T-cell exhaustion in e.g. chronic infection and cancer.

We also investigated at the whole transcriptome level how stem-like CD8^+^ T cells raised in our vaccination model compared to those raised by a tumor. For this purpose, we used a dataset derived from a genetically engineered mouse model of spontaneous lung cancer, wherein tumor cells express LCMV-derived foreign antigens^34^ (**Figure 5D**). The authors showed that tumor-specific PD-1^+^TCF-1^+^ stem-like cells accumulated in the tumor-draining LN (TdLN), and that these largely shared their transcriptome with stem-like CD8^+^ T cells from chronic LCMV infection. Module score analysis showed that the top-100 gene expression signature of TdLN-derived stem-like CD8^+^ T cells (**Supplementary Table 3**) was highly enriched in the stem-like CD8^+^ T cells raised in our vaccination model (**Figure 5E, F**). These data extrapolate our findings to a large array of data from commonly used mouse models of (chronic) infection and cancer, making it plausible that the TCF-1^+^ progenitor from which both effector and exhausted cells can arise equals a primed CD8^+^ T cell that has not (yet) received CD4^+^ T-cell help to differentiate into an effector CTL.

In an extension of the model of Pritykin *et al.* ^7^ and others^8,10^, based on our current findings and review of the literature, we propose the following model: During CD8^+^ T-cell priming, stem-like cells are generated prior to delivery of CD4^+^ T-cell help signals. In case help signals are subsequently delivered, these stem-like cells are driven along a progressive differentiation trajectory into fully functional effector CTLs. When help signals are lacking and stem-like CD8^+^ T cells are exposed to chronic antigen stimulation under conditions as found in chronic infection and cancer, they follow the differentiation trajectory towards terminal exhaustion (**Figure 5G**).

### The MC38 tumor primes neoantigen-specific stem-like, but not helped effector CD8^+^ T cells

We considered that CD4^+^ T-cell priming might be deficient in tumor settings, due to absence of helper epitopes and/or innate immune stimuli such as IFN-I^13,22^. As a result, tumors would prime helpless CD8^+^ T cells, which in continued absence of help signals and chronic presence of antigen become exhausted in the TME. To investigate this, we directly compared the differentiation states of CD8^+^ T cells of the same antigenic specificity that were either raised by Help vaccination or by the widely used immunogenic mouse colon carcinoma model MC38. A DNA vaccine was created that encodes defined MHC-I-restricted MC38 neoantigens derived from mutated Rpl18 and Adpgk^27^, in conjunction with the Help cassette (**Figure 6A**). Mice were either vaccinated with this DNA construct and analyzed at the peak of the response (day 10), or subcutaneously injected with MC38 cells at day 0, and analyzed at day 7 or 15 during tumor outgrowth (**Figure 6A**). Frequencies of Rpl18-tetramer^+^ CD8^+^ T cells raised by MC38 tumors at day 15 were similar to those raised by Help vaccination at day 10 in blood, spleen and dLN (**Figure 6B**). Rpl18-specific CD8^+^ T cells were also clearly detectable within the MC38 tumor (**Figure 6B**). We next compared neoantigen-specific CD8^+^ T-cell differentiation in the vaccination and tumor settings by flow cytometry (**Supplementary Figure 9A-C**). In tumor-bearing mice, the frequency of PD-1^+^TCF-1^+^SLAMF6^+^ stem-like cells within the Rpl18-specific CD8^+^ T-cell pool was significantly higher than in vaccinated mice, both dLN and spleen (**Figure 6C**). Conversely, in Help vaccinated mice, the frequency of KLRG1^+^CX3CR1^+^ effector cells within the Rpl18-specific CD8^+^ T-cell pool was significantly higher than in tumor-bearing mice (**Figure 6D**). These results indicate that the MC38 tumor predominantly primes stem-like CD8^+^ T cells and not help-dependent effector CTLs. Accordingly, the frequency of GZMB^+^ cells among Rpl18-specific CD8^+^ T cells and their GZMB protein levels were higher after vaccination than after MC38 tumor implantation (**Figure 6E-G**). Also, the frequency of IFNγ^+^TNFα^+^ cells within total CD8^+^ T cells was higher in vaccinated mice than in tumor-bearing mice, as revealed by *ex vivo* stimulation of splenocytes with Rpl18/Adpgk neoantigen peptides (**Figure 6H, I**). This reflected improved functional capacity rather than increased size of the responder population, since the frequency of antigen-specific CD8^+^ T cells in spleen was similar after vaccination or tumor inoculation (**Supplementary Figure 9D**).

**Figure 6.**
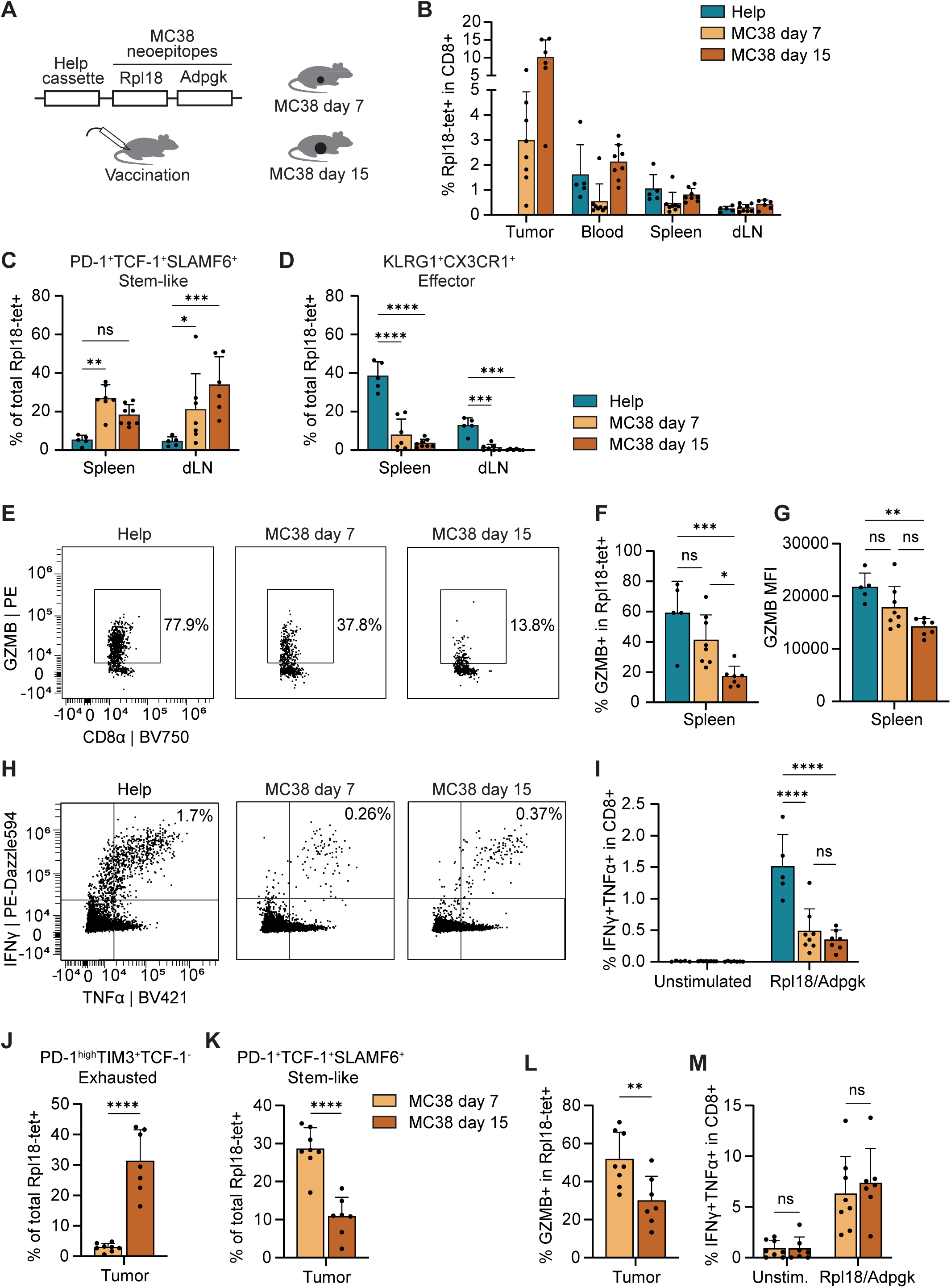
The MC38 tumor primes neoantigen-specific stem-like, but not helped effector CD8^+^ T cells. **(A)** Experimental setup. Mice were vaccinated on day 0, 3 and 6 with a DNA construct encoding the Help cassette and MC38-derived neo-epitopes from mutated Rpl18 and Adpgk^38^, or s.c. inoculated on day 0 with 300.000 MC38 tumor cells. CD8^+^ T-cell responses were analyzed by flow cytometry on day 10 after vaccination, or day 7 or 15 after tumor inoculation. (**B**) Frequency of Rpl18-tetramer^+^ CD8^+^ T cells in indicated tissues and the MC38 tumor. (**C, D**) Frequency of PD-1^+^TCF-1^+^ stem-like (C) and KLRG1^+^CX3CR1^+^ effector cells (D) within the Rpl18-specific CD8^+^ T-cell pool in the indicated organs. (**E-G**) Frequency of Granzyme-B (GZMB)^+^ cells within the Rpl18-specific CD8^+^ T cell population in spleen (E, F), and MFI of GZMB in this population (G). (**H, I**) Frequency of IFNγ^+^TNFα^+^ cells within the CD8^+^ T cell population from spleen after *ex vivo* stimulation with Rpl18/Adpgk neoantigen peptides. (**J, K**) Frequency of PD-1^hi^TIM3^+^TCF-1^-^ exhausted (J) and PD-1^+^TCF-1^+^ stem-like cells (K) within the CD8^+^ TIL pool on day 7 or 15 after tumor inoculation. (**L**) Frequency of GZMB^+^ cells within Rpl18-specific CD8^+^ TILs. (**M**) Frequency of IFNγ^+^TNFα^+^ cells within CD8^+^ TILs upon *ex vivo* restimulation with Rpl18/Adpgk peptides. Data in A-D and E-M are from 2 separate experiments with n=5-8 mice per group. Error bars indicate SD. Statistical significance was calculated by two-way (B-D, I, M) or one-way (F, G) ANOVA and Sidak’s multiple comparisons test, or unpaired t-test (J, K). *P < 0.05, **P < 0.005, ***P < 0.001 and ****P < 0.0001. See also Supplementary Figure 9.

We similarly examined the differentiation states of Rpl18-specific and total CD8^+^ T cells within the MC38 tumor (**Supplementary Figure 9E-I**). The great majority of Rpl18-specific CD8^+^ tumor infiltrating lymphocytes (TILs) expressed PD-1 and none of these cells had the KLRG1^+^CX3CR1^+^ helped effector phenotype, even though a fraction of them expressed GZMB (**Supplementary Figure 9F**). Also in the total CD8^+^ TIL population, GZMB^+^PD-1^+^ cells were found, while cells with a KLRG1^+^CX3CR1^+^ helped effector phenotype were lacking (**Supplementary Figure 9H, I**). Among Rpl18-specific TILs, the frequency of PD-1^high^TIM3^+^TCF-1^-^ exhausted cells^4^ increased from day 7 to day 15 (**Figure 6J**), while the population of stem-like PD-1^int^TCF-1^+^SLAMF6^+^ TILs decreased (**Figure 6K**), in line with the concept that stem-like cells become exhausted upon chronic antigen exposure in the TME. Accordingly, the frequency of Rpl18-specific GZMB^+^ TILs also decreased upon tumor progression (**Figure 6L**). Among total CD8^+^ TILs, a small proportion retained the capacity to produce IFNγ and TNFα after *in vitro* stimulation (**Figure 6M**).

We conclude that tumor-specific CD8^+^ T cells primed by MC38 reach a stem-like differentiation state that develops into exhaustion in the TME, while they do not reach an effector differentiation state as can be acquired after delivery of CD4^+^ T-cell help.

### Expression of vaccine helper epitopes in the MC38 tumor does not rescue helped effector differentiation of CD8^+^ TILs

MC38 tumor cells express MHC-I and MHC-II-restricted neoepitopes^27,29,44^, which in theory would enable priming of tumor-specific CD8^+^ and CD4^+^ T cells. However, the absence of a helped effector population in MC38 tumor-bearing mice suggests that even when tumor cells express CD4^+^ T-cell epitopes, help signals are not properly delivered. Potentially, endogenous neoepitopes are not immunogenic enough to prime a CD4^+^ T-helper response, or DC-mediated help delivery from CD4^+^ to CD8^+^ T cells is impaired in the tumor setting. To increase the likelihood of help delivery, we stably expressed in wild-type (wt) MC38 cells the strong helper epitopes P30 and PADRE as present in our vaccine HELP cassette (**Figure 7A**). Next, we compared tumor-specific CD8^+^ T-cell responses and tumor outgrowth in the MC38 wt and MC38-HELP settings. The exogenous helper epitopes in the MC38-HELP tumor were immunogenic as shown by the presence of a FOXP3^-^ CD44^+^PADRE-tetramer^+^ CD4^+^ T-cell population in the spleen of mice bearing the MC38-HELP tumor, but not the MC38 wt tumor (**Figure 7B, Supplementary Figure 10A**). Outgrowth of MC38-HELP tumors was less efficient than that of MC38 wt tumors, resulting in lower tumor weight by day 15 and tumor clearance in 2 out of 10 mice (**Figure 7C, D**). The frequency of Rpl18-specific cells among CD8^+^ TILs was higher in MC38-HELP than in MC38 wt by day 15 (**Figure 7E**). However, the frequency of KLRG1^+^CX3CR1^+^ helped effector cells among Rpl18-tetramer^+^ CD8^+^ TILs was not different between the tumor settings (**Figure 7F**). Overall, the differentiation state of Rpl18-specific CD8^+^ T cells in MC38 wt and MC38-HELP tumors did not differ as assessed by the panel of differentiation markers, and most cells expressed PD-1 (**Supplementary Figure 10B-E**). The frequency of GZMB^+^ cells among Rpl18-tetramer^+^ CD8^+^ TILs and total CD8^+^ TILs was also not different between the two tumor settings (**Figure 7G, H**) and these were of a KLRG1^-^CX3CR1^-^PD-1^+^ phenotype (**Supplementary Figure 10E-H**). CD8^+^ TILs in both tumor settings had similar phenotypes, were largely PD-1^+^ and cells with a KLRG1^+^CX3CR1^+^ helped effector phenotype were lacking. Likewise, the frequencies of CD8^+^ TILs capable of producing IFNγ and TNFα upon *ex vivo* stimulation with Rpl18/Adpgk neoantigen peptides did not differ (**Figure 7I**). These results indicate that the expression of strong helper epitopes by MC38 tumors can raise tumor-specific CD4^+^ T cells, but these do not rescue the formation of tumor-specific KLRG1^+^CX3CR1^+^ helped effector CD8^+^ TILs, suggesting that the delivery of CD4^+^ T-cell help is impaired in this tumor context.

**Figure 7.**
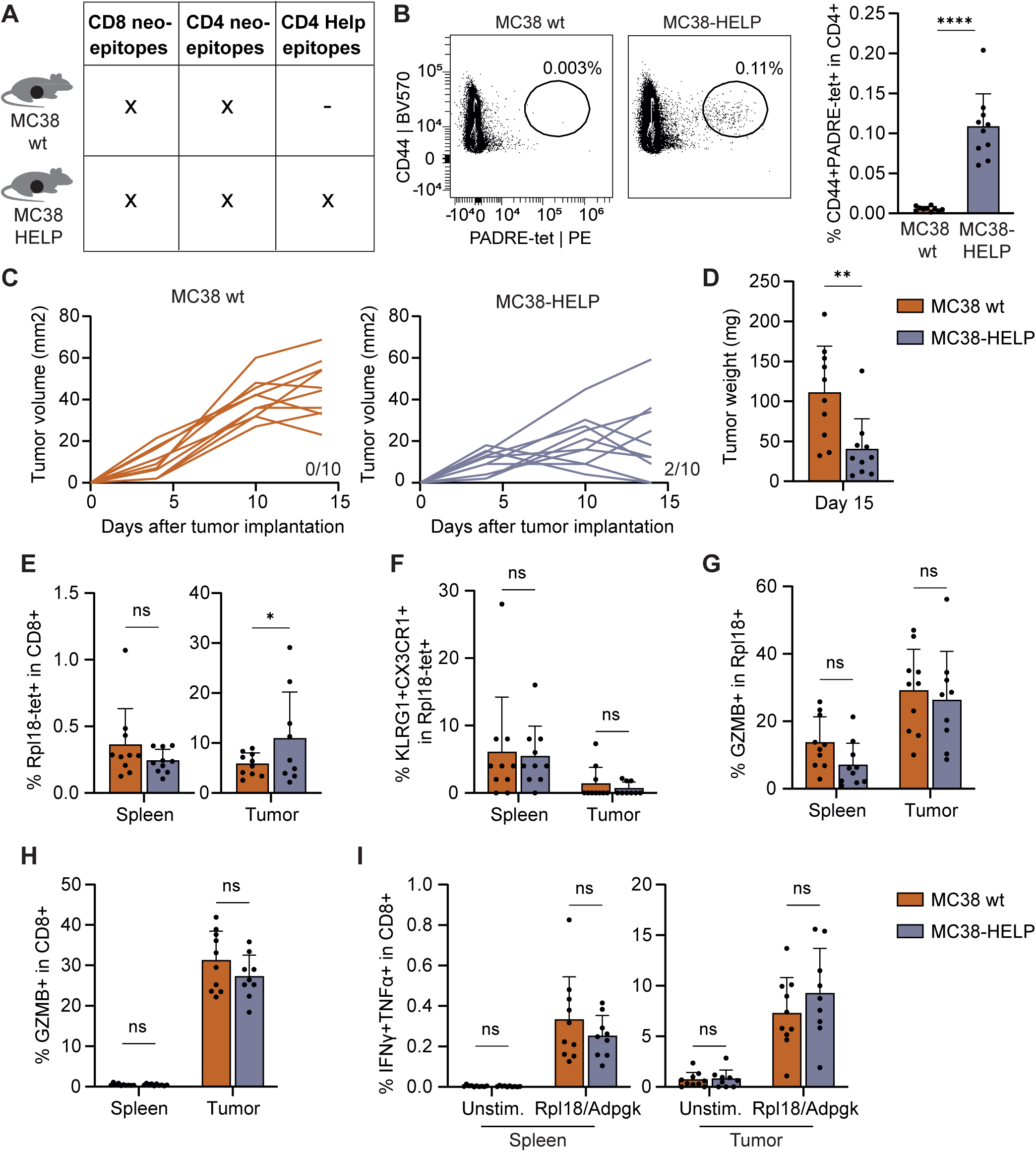
Expression of vaccine helper epitopes by MC38 tumors does not rescue helped effector differentiation of CD8^+^ TILs. **(A)** MC38 tumor cells for which endogenous MHC-I and MHC-II restricted neoepitopes have been defined were transduced to stably express the Help cassette (P30/PADRE) as present in our vaccine, resulting in the MC38-HELP tumor line. (**B-I**) Mice were inoculated with 300.000 MC38 or MC38-HELP cells (legend for all panels as indicated in Panels D and I) at day 0, and analyzed at day 15. (**B**) Frequency of CD44^+^PADRE-tetramer^+^ among CD4^+^ T cells in spleen. (**C**) Outgrowth of MC38 and MC38-HELP tumors until day 15. Numbers of mice with complete tumor clearance are indicated. (**D**) Tumor weight at day 15. (**E**) Frequency of Rpl18-tetramer^+^ CD8^+^ T cells in spleen and tumor. (**F**) Frequency of KLRG1^+^CX3CR1^+^ effector cells within the Rpl18-specific CD8^+^ T-cell pool in the indicated organs. (**G**) Frequency of Granzyme-B (GZMB)^+^ cells within Rpl18-specific CD8^+^ T cells in spleen and tumor. (**H**) Frequency of GZMB^+^ cells within total CD8^+^ T cells in spleen and tumor. (**I**) Frequency of IFNγ^+^TNFα^+^ cells within CD8^+^ T cells from spleen and tumor upon *ex vivo* restimulation with Rpl18/Adpgk neoantigen peptides. Data are from 1 experiment representative of 2 independent experiments with n=10 mice per group. Error bars indicate SD. Statistical significance was calculated by two-way ANOVA and Sidak’s multiple comparisons test (E-H), or unpaired t-test (B,D). *P < 0.05, **P < 0.005, ***P < 0.001 and ****P < 0.0001. See also Supplementary Figure 10.

## DISCUSSION

Here, we demonstrate that during priming, CD4^+^ T-cell help shifts the CD8^+^ T-cell differentiation spectrum beyond the stem-like state to the CTL effector state. Our data show that cytotoxic and migratory qualities of CD8^+^ T cells are installed during their differentiation from stem-like to effector CTLs, which is dependent on CD4^+^ T-cell help. During this phase, antigen-specific CD8^+^ T cells also undergo clonal expansion. The data explain the increased functionality that has been described for helped CD8^+^ T cells^15–17^ as a shift in the differentiation spectrum of the primed CD8^+^ T-cell population from stem-like to more effector-like. Importantly, we did not exclusively raise effector CTLs upon Help vaccination, but cells ranging along the entire CD8^+^ T-cell differentiation spectrum, including stem-like cells. On phenotypic and transcriptomic level, stem-like cells from the Help and No Help vaccination settings were identical, suggesting that these cells have received the same signals during priming. We propose therefore that stem-like cells generated after Help vaccination also represent helpless cells, that have undergone the first step of priming but have not yet received CD4^+^ T-cell help. After Help vaccination, stem-like cells in the dLN turn into effector cells over time, in accordance with the idea that help is delivered in a second step of priming.

In the progressive differentiation model, CD8^+^ T-cell fate has been proposed to be mainly determined by TCR signaling strength and inflammatory signals during priming^1,2^. We specify here that effector differentiation additionally depends on CD4^+^ T-cell help that allows cDC1s to provide specific cytokine- and costimulatory signals to CD8^+^ T cells^12–14^. We propose that the final differentiation fate of CD8^+^ T cells is determined by the integration of TCR-, costimulatory and cytokine signals received during both the first and the second priming steps. This would also explain why CTL responses are differentially dependent on CD4^+^ T-cell help in diverse infection, tumor or vaccination settings, that vary in the amount of antigen and inflammatory signals they elicit^21^. Our findings support the concept that CD8^+^ stem-like cells lie on the trajectory towards effector differentiation, but can alternatively differentiate towards terminal exhaustion such as in cancer and chronic infection, as recently suggested^7,10^. Exhaustion depends on persistent antigen exposure of the stem-like population^3,6^, but in our vaccination setting antigen is rapidly lost due to turnover of transfected keratinocytes^35,36^. Accordingly, we did not observe exhaustion of CD8^+^ T cells as defined by a PD-1^high^TIM3^+^TCF-1^-^ phenotype after vaccination.

Delivering CD4^+^ T-cell help signals after priming allowed stem-like CD8^+^ T cells to further proliferate and complete their CTL effector differentiation in our vaccination model. Ideally, a similar approach may improve the CD8^+^ T-cell response under more challenging conditions such as chronic infection and cancer. In chronic LCMV infection, proliferative and cytotoxic capacities of stem-like CD8^+^ T cells were improved by adoptive transfer of virus-specific CD4^+^ T cells^23,24^. Adoptive transfer of neoantigen-specific CD4^+^ T cells or expression of MHC-II-restricted neoantigens by tumor cells have been shown to increase CD8^+^ T cell-mediated cytotoxicity and tumor control in mouse tumor models^45,46^. However, we found that expression of strong helper epitopes by MC38 tumor cells raised an antigen-specific CD4^+^ T-cell response but still failed to induce KLRG1^+^CX3CR1^+^ helped effector CTLs. These data illustrate that presence of helper epitopes in a tumor does not guarantee that help signals are actually delivered to CD8^+^ T cells. We did observe improved control of the MHC-II negative MC38 tumor^29^, which may be attributed to other functions of tumor-specific CD4^+^ T cells, such as cytokine production or help for B cells^47^. TMEs of different tumor types are highly diverse and various circumstances may prevent help delivery, for example lack of inflammatory signals such as IFN-I^22^ or the induction of Treg responses that impede effective CD8^+^ T-cell priming^48,49^.

The validity of our CD8^+^ T-cell differentiation model should be further investigated in other settings, such as viral infection and human cancer. Nevertheless, we propose that helpless priming may be causally related to CD8^+^ T-cell exhaustion, since lack of helped effector CTLs will lead to antigen persistence^11^. We hope that our study contributes to a theoretical framework on CTL effector differentiation that provides a rationale to optimize immunotherapeutic strategies.

## DECLARATIONS

## Acknowledgements

We thank Kees Franken for manufacturing tetramers, Mart de Boo for helping with tissue processing, Fiamma Salerno and Elselien Frijlink for scientific discussions, Ferry Ossendorp for advice on the tumor model, Susan Kloet for advise on scRNAseq, Tom de Wit for help with sorting, Evert de Vries for general support, Nikhil Yoshi and Kelli Connolly for sharing their dataset, and the Flow Cytometry Facility, Leiden Genome Technology Center, and Experimental Animal Facility of our institute for technical assistance.

## Author contributions

Study design: J.Bo, J.Bu; experimental design and performing experiments: J.Bu, D.B., M.S., Y.X, X.L; data analysis: J.Bu; manufacturing DNA vaccines: M.S.; paper writing: J.Bu, J.Bo; critical reading and editing of the paper: D.B., M.S., X.L, Y.X.

## LIST OF ABBREVIATIONS

cDC: conventional dendritic cell
CTL: cytotoxic T lymphocyte
DC: dendritic cell
dLN: draining lymph node
HPV: human papilloma virus
LCMV: lymphocytic choriomeningitis virus
LN: lymph node
ndLN: non-draining lymph node
OVA: ovalbumin
scRNAseq: single cell RNA sequencing
T_CM_: T central memory
TCR: T cell receptor
TCRseq: T cell receptor sequencing
TdLN: tumor-draining lymph node
T_EM_: T effector memory
tet: tetramer
TIL: tumor-infiltrating lymphocyte
TME: tumor micro-environment
T_SCM_: T stem cell memory.

