## Supplementary Figures for "Stem-like PD-1^+^TCF-1^+^ CD8^+^ T cells result from helpless priming and rely on CD4^+^ T-cell help to complete their cytotoxic effector differentiation"

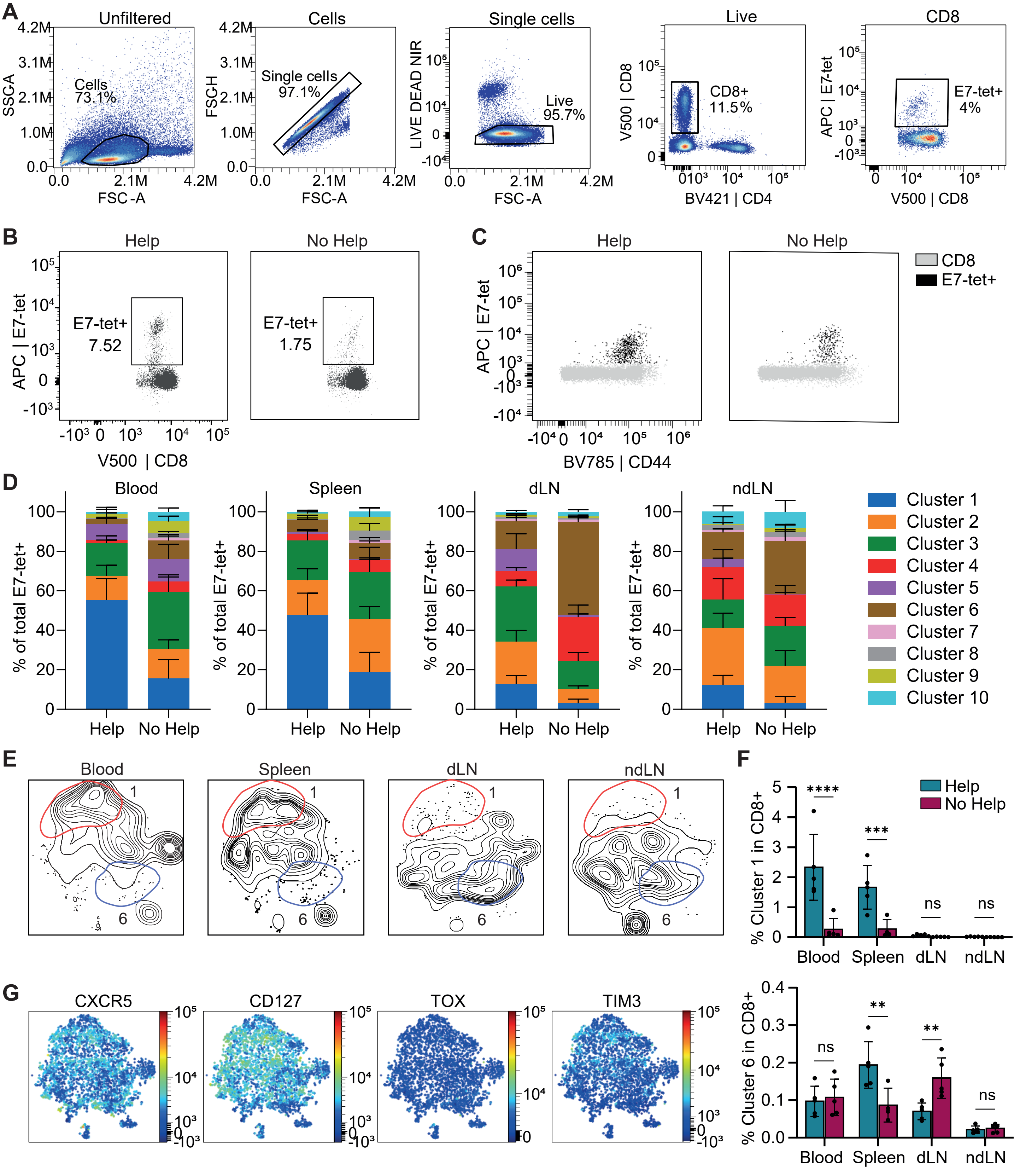


**Supplementary Figure 1. Gating and phenotyping of antigen-specific CD8⁺ T cells after Help and No Help vaccination**

Mice were vaccinated under Help or No Help conditions at day 0, 3, and 6 and sacrificed at day 10 after which blood, spleen, dLN and ndLN were analyzed as described in Figure 1. (**A**) Gating strategy for antigen-specific (E7-tetramer⁺) CD8⁺ T cells at day 10 after vaccination. Gating strategy is shown for 1 blood sample, but was used consistently for all experiments and organs. (**B-C**) Representative FACS plots from one mouse per group, showing E7-tetramer⁺ CD8⁺ cells in blood at day 11 after vaccination (B) and CD44 expression in E7-tet⁺ and total CD8⁺ in spleen at day 10 (C). (**D-G**) Opt-SNE visualization and FlowSom clustering of E7-tetramer⁺ CD8⁺ T cells at day 10 after Help or No Help vaccination was performed as shown in Figure 1E. (**D**) Frequency of tetramer⁺ CD8⁺ T cells per cluster per organ. (**E**) Distribution of cells from different organs across opt-SNE. (**F**) Frequency of Cluster 1 and Cluster 6 E7-tetramer⁺ cells in total CD8⁺ T cells per organ after Help versus No Help vaccination. (**G**) Expression of additional differentiation markers on cumulative E7-tetramer⁺ CD8⁺ T cells from all experimental settings visualized by opt-SNE. Data are from 1 experiment, representative of 2 independent experiments with n=5 mice per group.

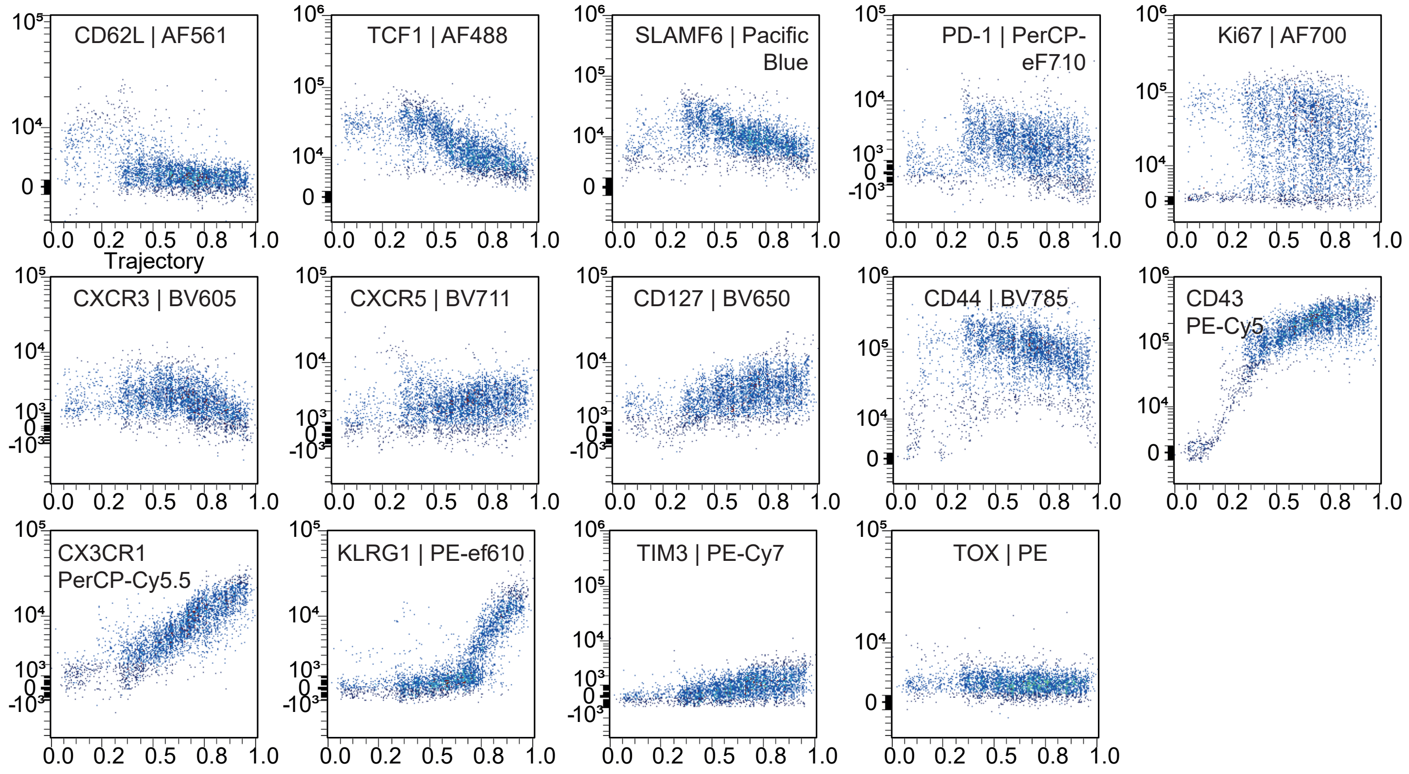


**Supplementary Figure 2. Expression of marker proteins along the CTL differentiation spectrum upon vaccination**

Antigen-specific E7-tetramer$^{+}$ CD8$^{+}$ T cells from blood, spleen, dLN and ndLN at day 10 after Help or No Help vaccination were analyzed by flow cytometry using the differentiation marker panel, as shown in Figure 2. FACS plots showing expression levels of each differentiation marker against the Wanderlust trajectory score. Data are from 1 experiment, representative of 2 independent experiments with n=5 mice per group.

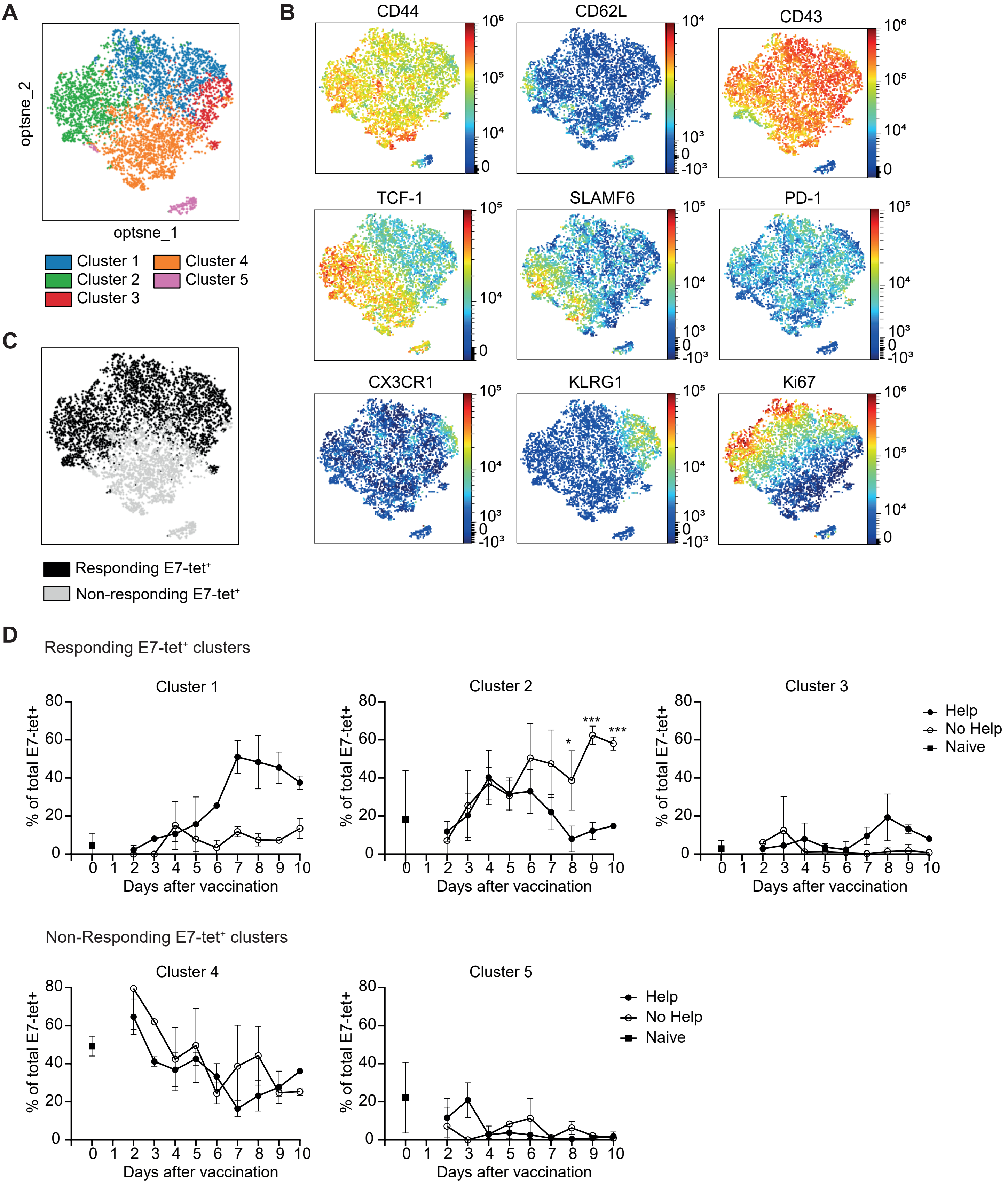


**Supplementary Figure 3. Selection of responding E7-tetramer⁺ clusters for analysis over time**

Experimental details related to the timecourse experiment depicted in Figure 3. Mice received Help or No Help vaccination at day 0, 3 and 6, and different cohorts of mice were sacrificed at day 2-10 after first vaccination (**A**) Opt-SNE dimension reduction and FlowSom clustering of E7-tetramer⁺ CD8⁺ T cells form dLN of vaccinated mice (n=3 per group per timepoint), and 3 naive LNs. (**B**) Expression of differentiation markers on cumulative E7-tetramer⁺ CD8⁺ T cells from all experimental settings visualized by opt-SNE. (**C**) Responding and non-responding E7-tetramer⁺ CD8⁺ T cells in opt-SNE. (**D**) Frequencies of responding E7-tetramer⁺ clusters and non-responding E7-tetramer⁺ clusters in naive LNs and at different timepoints after vaccination. Responding clusters were determined by a frequency of <20% in naive LN, and a change in frequency between different timepoints after vaccination. Responding E7-tetramer⁺ CD8⁺ T cells from vaccinated mice (day 2-10) were used for further analysis as shown in Figure 3.

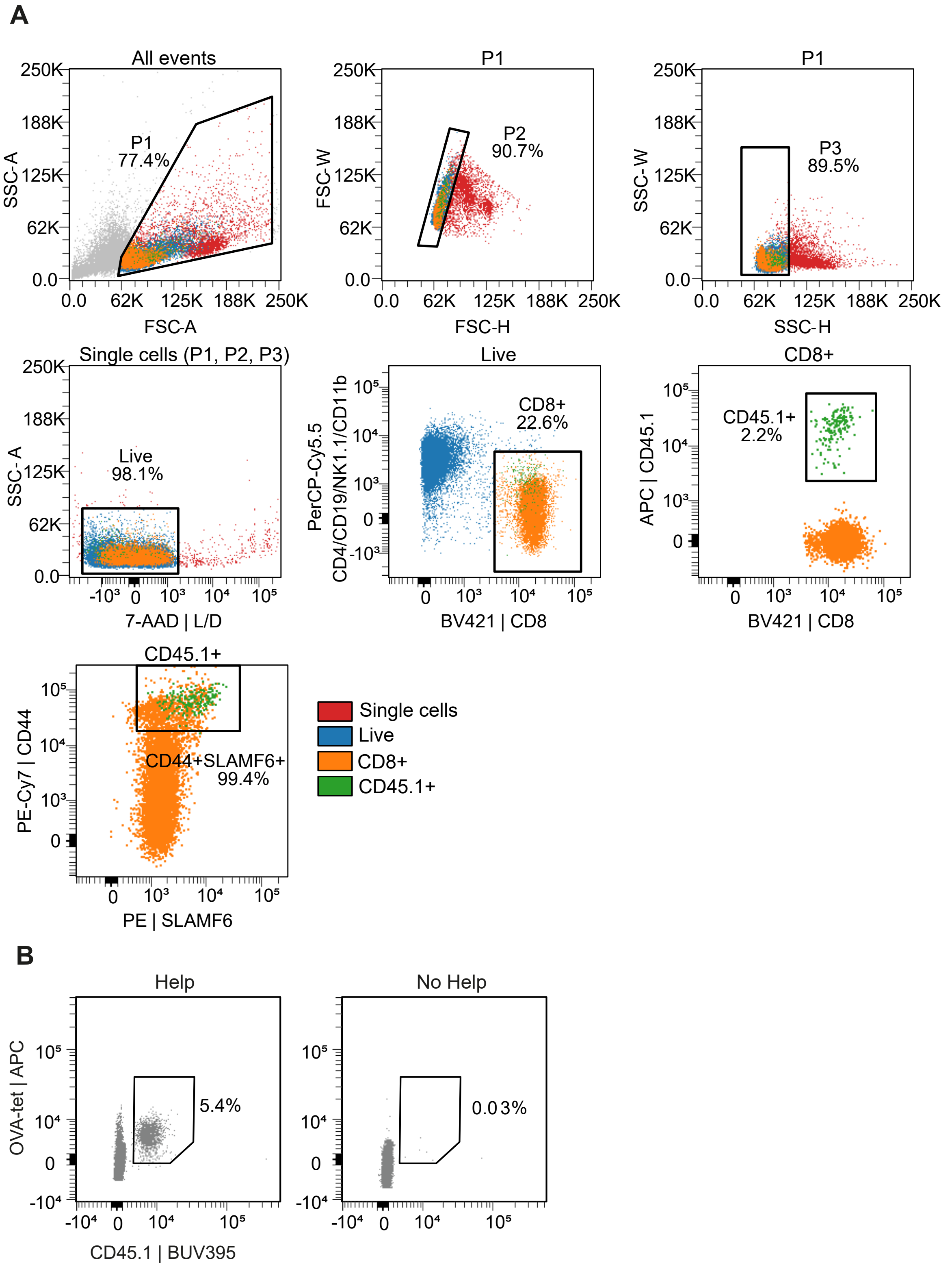


**Supplementary Figure 4. Adoptive transfer of helpless OT-I cells**

Naïve CD45.1$^+$ OVA$_{257/264}$/H-2K$^b$-specific OT-I cells were adoptively transferred to CD45.2$^+$ recipient mice, which were subsequently vaccinated with No Help vaccine encoding the OVA$_{257/264}$ epitope, as shown in Figure 3. (**A**) Sorting strategy of CD44$^+$SLAMF6$^+$CD45.1$^+$CD8$^+$ (OT-I) T cells from dLN at day 10 after vaccination of the primary recipeint mice. Sorting was performed on pooled dLNs from 5 mice and these cells were used for adoptive transfer into secondary CD45.2$^+$ recipient mice. (**B**) Representative FACS plots from one mouse per group, showing OT-I cells in blood at day 21, which is day 10 after secondary recipient vaccination.

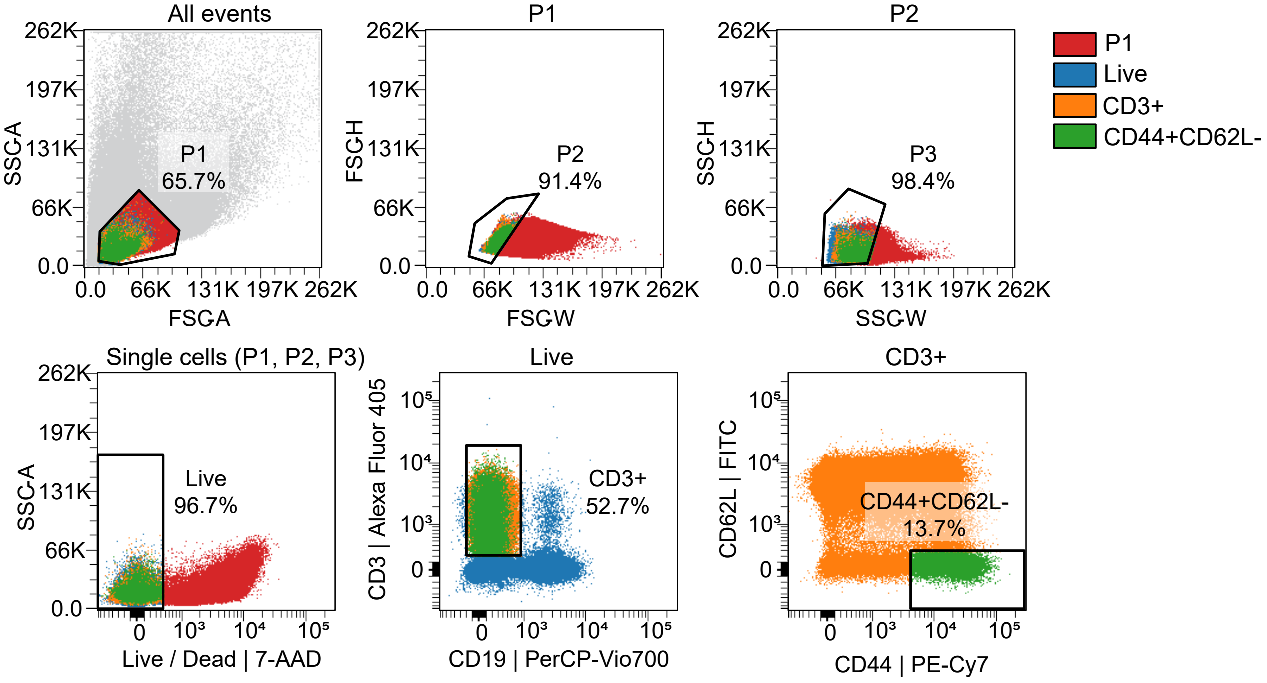


**Supplementary Figure 5. Sorting strategy of activated T cells for scRNAseq**

Sorting strategy for the scRNAseq experiment depicted in Figure 4. Mice were vaccinated under E7-Help or No Help conditions (n=3 per group) and activated T cells were flow cytometrically sorted per experimental group from pooled dLN or ndLN at day 5 or day 10, on a $CD3^{+}CD44^{+}CD62L^{-}$ phenotype.

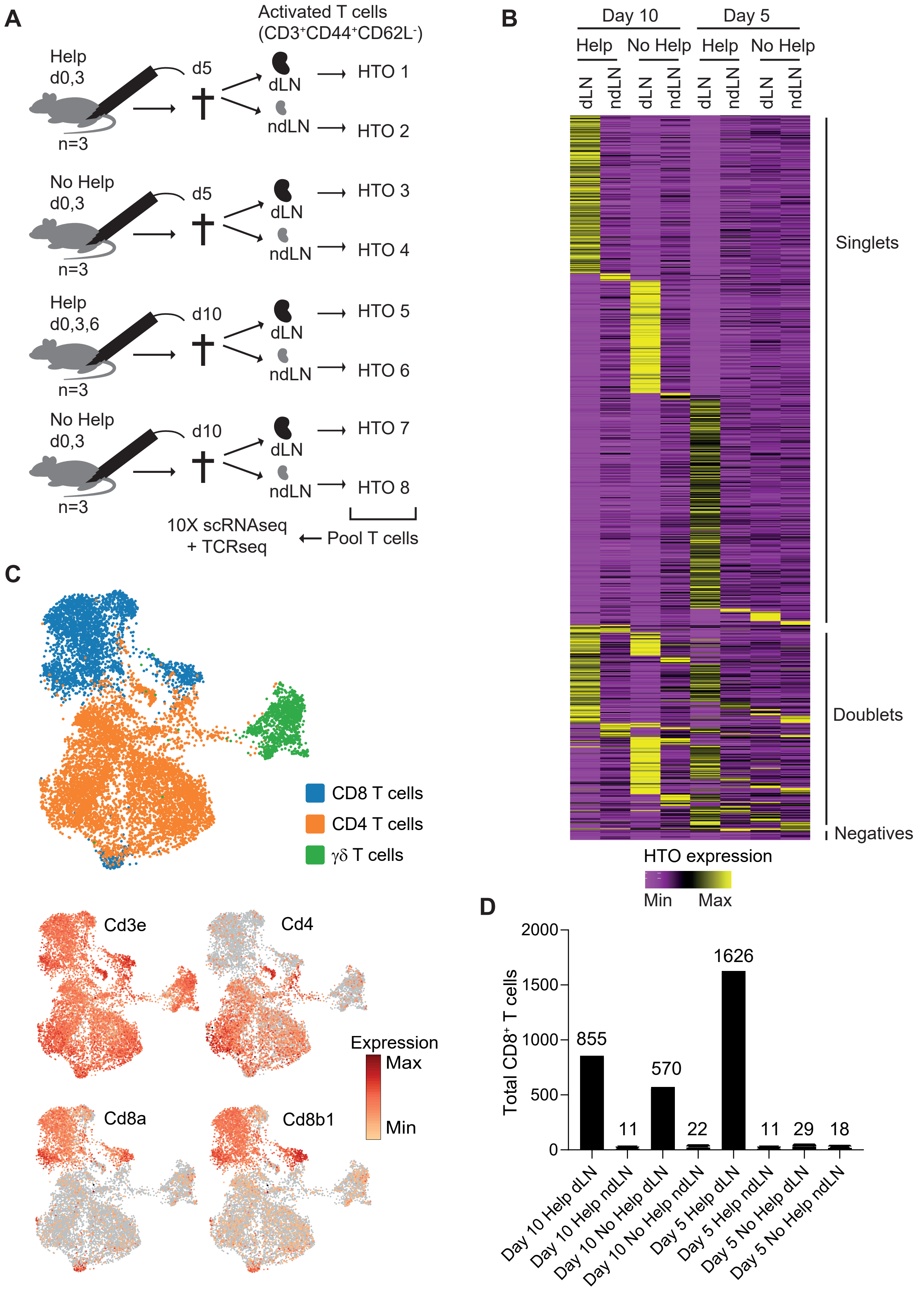


**Supplementary Figure 6. Generation and processing of scRNAseq data of CD8$^+$ T cells after Help or No Help vaccination**

Experimental details related to the scRNAseq experiment depicted in Figure 4. (**A**) Experimental setup. Mice (n=3 per group) received E7-Help or No Help vaccination at the indicated days and were sacrificed at day 5 or day 10, after which dLN and ndLN were isolated and pooled per group. Activated T cells were sorted and labeled with MHC-II/CD45 antibody-coupled hashtag oligonucleotide (HTO) as indicated. T cells from all HTO groups were pooled and subjected to 10x Genomics scRNAseq, coupled to TCRseq. (**B**) Heatmap showing HTO demultiplexing results. The color scale indicates expression levels of HTO corresponding to each experimental condition per cell. (**C**) Expression of canonical T cell marker genes across UMAP of activated T cells, for identification of CD8$^+$, CD4$^+$ and γδT cells. (**D**) Number of CD8$^+$ T cells per experimental condition after the entire procedure, as identified by HTO expression level.

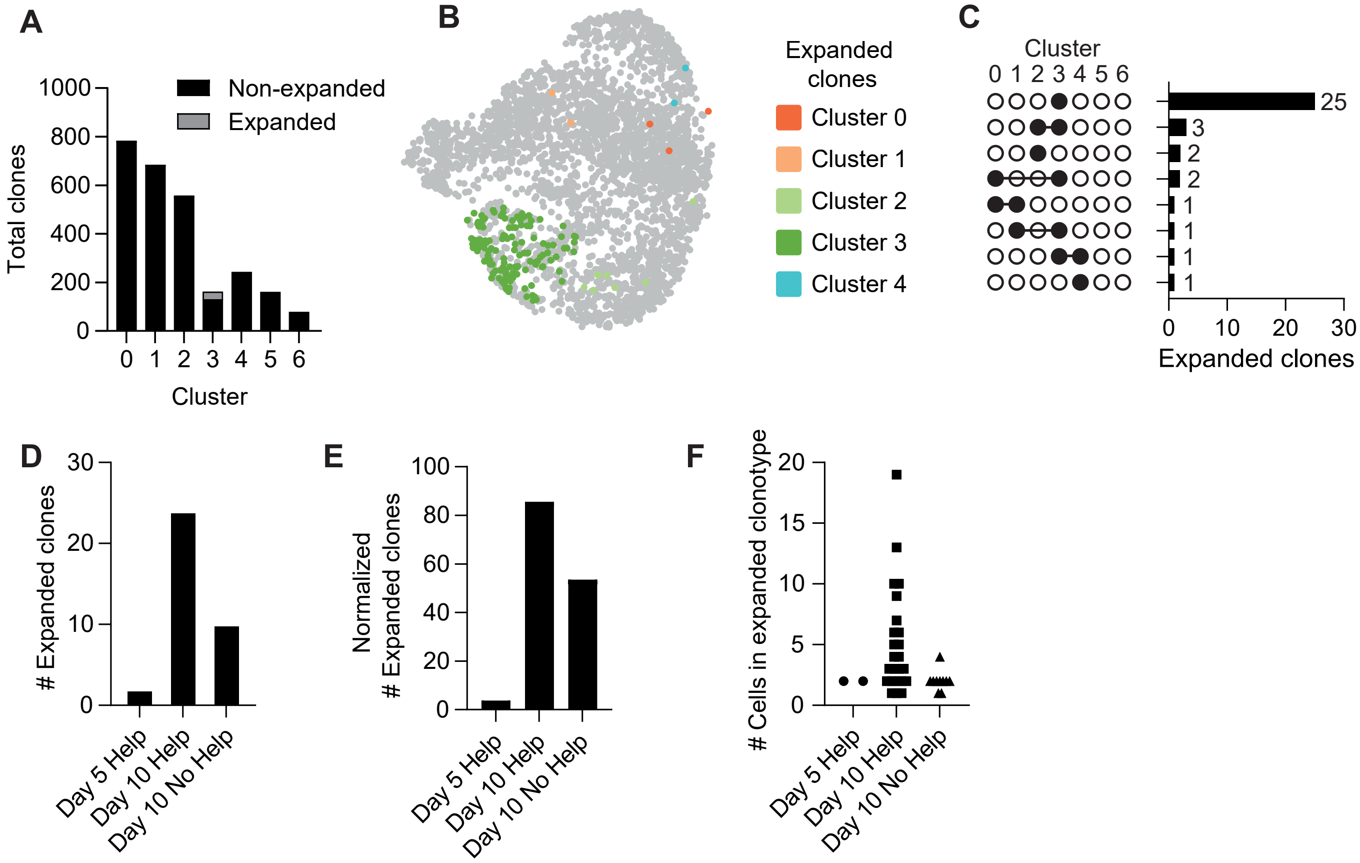


**Supplementary Figure 7. Quantification and characterization of expanded TCR clones within CD8⁺ T cells after vaccination under Help or No Help conditions**

Experimental details related to the TCRseq analysis depicted in Figure 4. (**A**) Quantification of distinct TCR clones that are non-expanded (containing 1 cell) or expanded (containing 2 or more cells) in each of the $CD8^+$ T cell clusters identified in Figure 4A. (**B**) UMAP showing $CD8^+$ T cells from expanded clones per $CD8^+$ T cell cluster. (**C**) Number of unique expanded TCR clones, of which all cells are found within one cluster (indicated by one black circle), and shared TCR clones, containing cells that are found in multiple clusters (indicated by multiple connected black circles). (**D, E**) Number of expanded TCR clones per experimental condition, absolute (D) and normalized to number of cells per condition (E). (**F**) Number of cells belonging to each expanded clonotype per experimental condition.

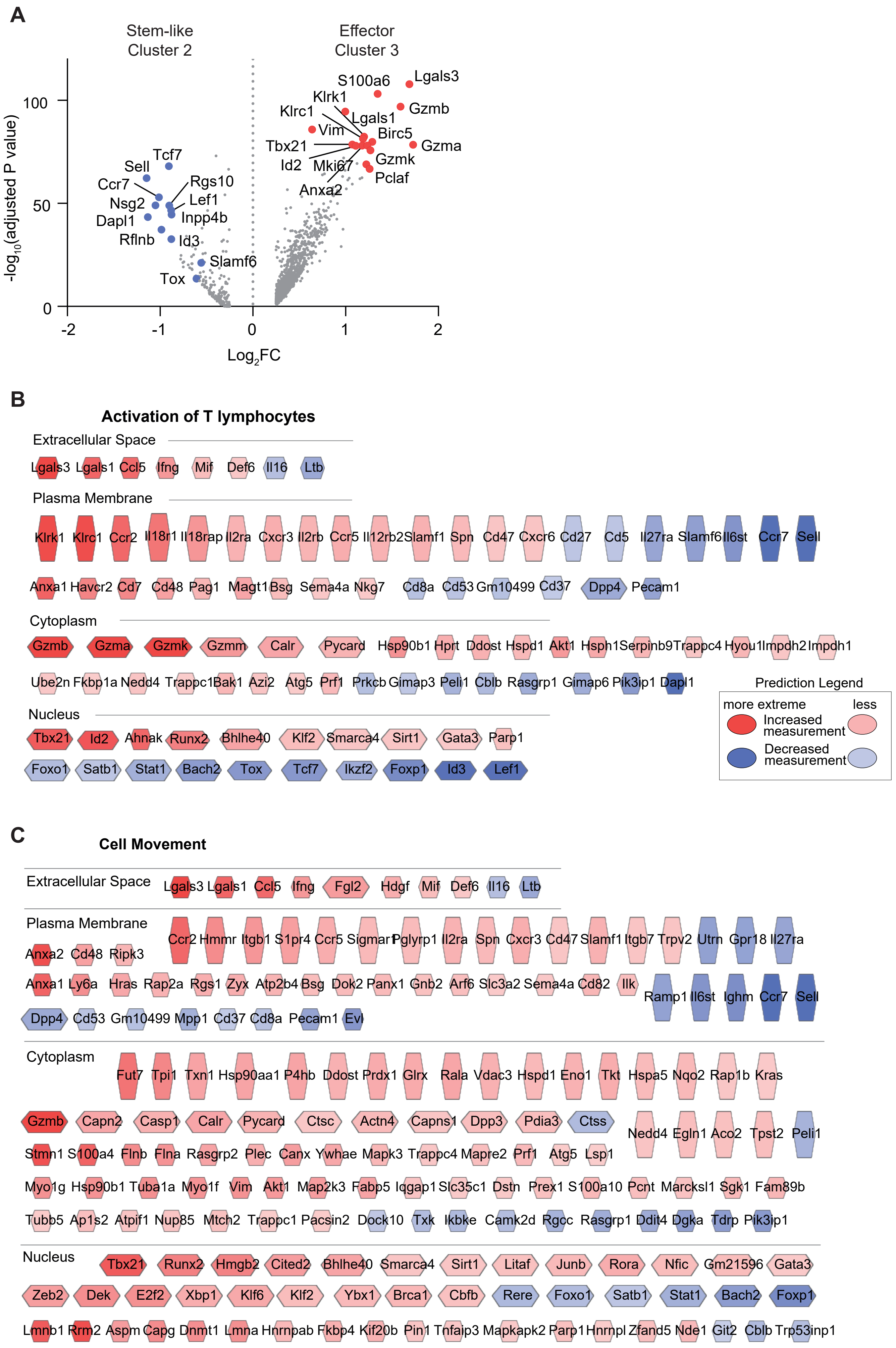
A
Stem-like Cluster 2
Effector Cluster 3
-log10(adjusted P value)
Log2FC
100
50
0
-2
-1
0
1
2
Tcf7
Sell
Ccr7
Nsg2
Dapl1
Rflnb
Rgs10
Lef1
Inpp4b
Id3
Slamf6
Tox
S100a6
Lgals3
Klrk1
Klrc1
Gzmb
Lgals1
Vim
Birc5
Tbx21
Gzma
Id2
Mki67
Gzmk
Pclaf
Anxa2
B
Activation of T lymphocytes
Extracellular Space
Lgals3 Lgals1 Ccl5 Ifng Mif Def6 Il16 Ltb
Plasma Membrane
Klrk1 Klrc1 Ccr2 Il18r1 Il18rap Il2ra Cxcr3 Il2rb Ccr5 Il12rb2 Slamf1 Spn Cd47 Cxcr6 Cd27 Cd5 Il27ra Slamf6 Il6st Ccr7 Sell
Anxa1 Havcr2 Cd7 Cd48 Pag1 Magt1 Bsg Sema4a Nkg7 Cd8a Cd53 Gm10499 Cd37 Dpp4 Pecam1
Cytoplasm
Gzmb Gzma Gzmk Gzmm Calr Pycard Hsp90b1 Hprt Ddost Hspd1 Akt1 Hsph1 Serpinb9 Trappc4 Hyou1 Impdh2 Impdh1
Ube2n Fkbp1a Nedd4 Trappc1 Bak1 Azi2 Atg5 Prf1 Prkcb Gimap3 Peli1 Cblb Rasgrp1 Gimap6 Pik3ip1 Dapl1
Nucleus
Tbx21 Id2 Ahnak Runx2 Bhlhe40 Klf2 Smarca4 Sirt1 Gata3 Parp1
Foxo1 Satb1 Stat1 Bach2 Tox Tcf7 Ikzf2 Foxp1 Id3 Lef1
Prediction Legend
more extreme
less
Increased measurement
Decreased measurement
C
Cell Movement
Extracellular Space
Lgals3 Lgals1 Ccl5 Ifng Fgl2 Hdgf Mif Def6 Il16 Ltb
Plasma Membrane
Ccr2 Hmmr Itgb1 S1pr4 Ccr5 Sigmar1 Pglyrp1 Il2ra Spn Cxcr3 Cd47 Slamf1 Itgb7 Trpv2 Utrn Gpr18 Il27ra
Anxa2 Cd48 Ripk3
Anxa1 Ly6a Hras Rap2a Rgs1 Zyx Atp2b4 Bsg Dok2 Panx1 Gnb2 Arf6 Slc3a2 Sema4a Cd82 Ilk Ramp1 Il6st Ighm Ccr7 Sell
Dpp4 Cd53 Gm10499 Mpp1 Cd37 Cd8a Pecam1 Evl
Cytoplasm
Fut7 Tpi1 Txn1 Hsp90aa1 P4hb Ddost Prdx1 Glrx Rala Vdac3 Hspd1 Eno1 Tkt Hspa5 Nqo2 Rap1b Kras
Gzmb Capn2 Casp1 Calr Pycard Ctsc Actn4 Capns1 Dpp3 Pdia3 Ctss Nedd4 Egln1 Aco2 Tpst2 Peli1
Stmn1 S100a4 Flnb Flna Rasgrp2 Plec Canx Ywhae Mapk3 Trappc4 Mapre2 Prf1 Atg5 Lsp1
Myo1g Hsp90b1 Tuba1a Myo1f Vim Akt1 Map2k3 Fabp5 Iqgap1 Slc35c1 Dstn Prex1 S100a10 Pcnt Marcksl1 Sgk1 Fam89b
Tubb5 Ap1s2 Atpif1 Nup85 Mtch2 Trappc1 Pacsin2 Dock10 Txk Ikbke Camk2d Rgcc Rasgrp1 Ddit4 Dgka Tdrp Pik3ip1
Nucleus
Tbx21 Runx2 Hmgb2 Cited2 Bhlhe40 Smarca4 Sirt1 Litaf Junb Rora Nfic Gm21596 Gata3
Zeb2 Dek E2f2 Xbp1 Klf6 Klf2 Ybx1 Brca1 Cbfb Rere Foxo1 Satb1 Stat1 Bach2 Foxp1
Lmnb1 Rrm2 Aspm Capg Dnmt1 Lmna Hnrnpab Fkbp4 Kif20b Pin1 Tnfaip3 Mapkapk2 Parp1 Hnrnpl Zfand5 Nde1 Git2 Cblb Trp53inp1

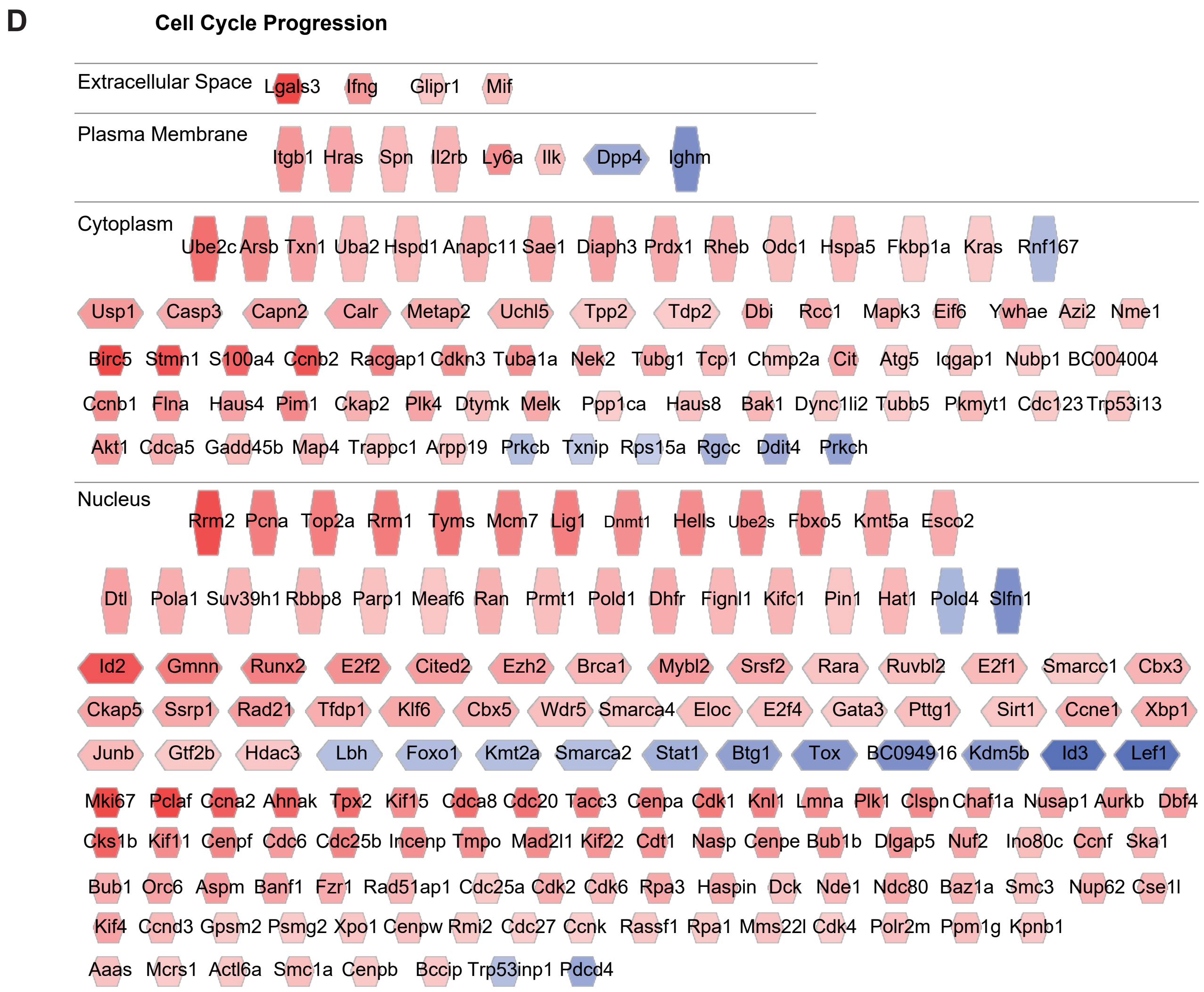


**Supplementary Figure 8. Functional differences between effector and stem-like CD8$^{+}$ T cells**

(**A**) Volcano plot showing genes that are differentially expressed between Cluster 2 and Cluster 3. (**B-D**) Ingenuity Pathway Analysis (IPA) showing transcripts in the category “Activation of T lymphocytes” (B), “Cell Movement” (C) and “Cell Cycle Progression” (D), with high (red) or low (blue) expression levels in helped effector Cluster 3 as compared to stem-like Cluster 2.

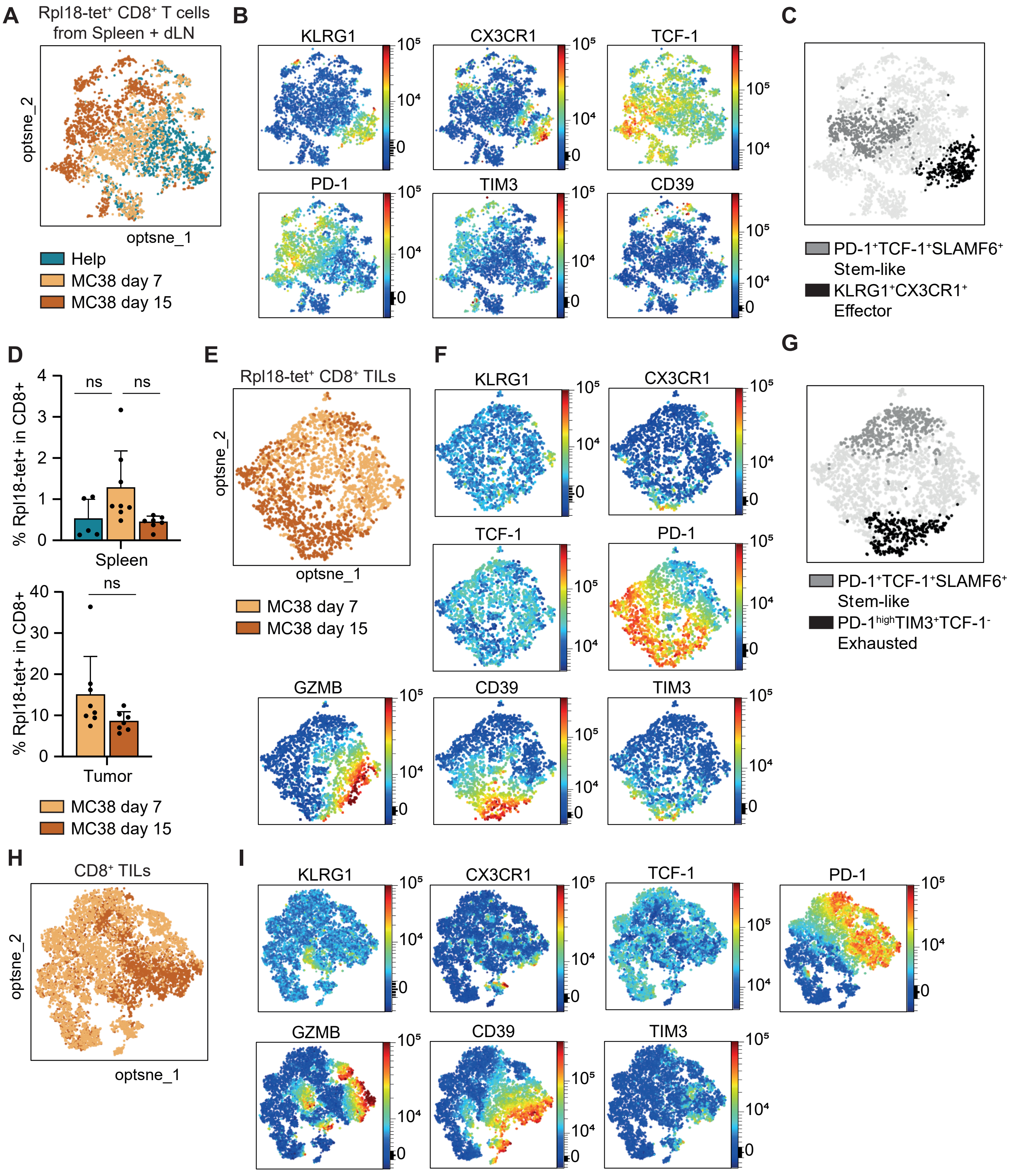


**Supplementary Figure 9. Characterization of antigen-specific CD8$^+$ T cells raised after vaccination or tumor implantation.** (**A**) Opt-SNE visualization of Rpl18-tetramer$^+$ CD8$^+$ T cells from spleen and dLN after Help vaccination or MC38 tumor implantation, as described in Figure 6. (**B**) Expression of differentiation markers on cumulative Rpl18-tetramer$^+$ CD8$^+$ T cells in spleen and dLN from the experimental settings visualized by opt-SNE in panel A. (**C**) Visualization of the PD-1$^+$TCF-1$^+$ stem-like and KLRG1$^+$CX3CR1$^+$ effector populations within opt-SNE. (**D**) Percentage of Rpl18-tetramer$^+$ CD8$^+$ T cells in spleen and the MC38 tumor, in the experiment depicted in Figure 7E-M. (**E**) Opt-SNE visualization of Rpl18-tetramer$^+$ CD8$^+$ TILs harvested from MC38 tumors at days 7 or day 15. (**F**) Expression of differentiation markers on cumulative Rpl18-tetramer$^+$ CD8$^+$ TILs visualized by opt-SNE. (**G**) Visualization of the PD-1$^{hi}$TIM3$^+$TCF-1$^-$ exhausted and PD-1$^+$TCF-1$^+$ stem-like populations within opt-SNE. (**H**) Opt-SNE visualization of total CD8$^+$ TILs harvested from MC38 tumors at days 7 or day 15. (**I**) Expression of differentiation markers on cumulative total CD8$^+$ TILs visualized by opt-SNE.

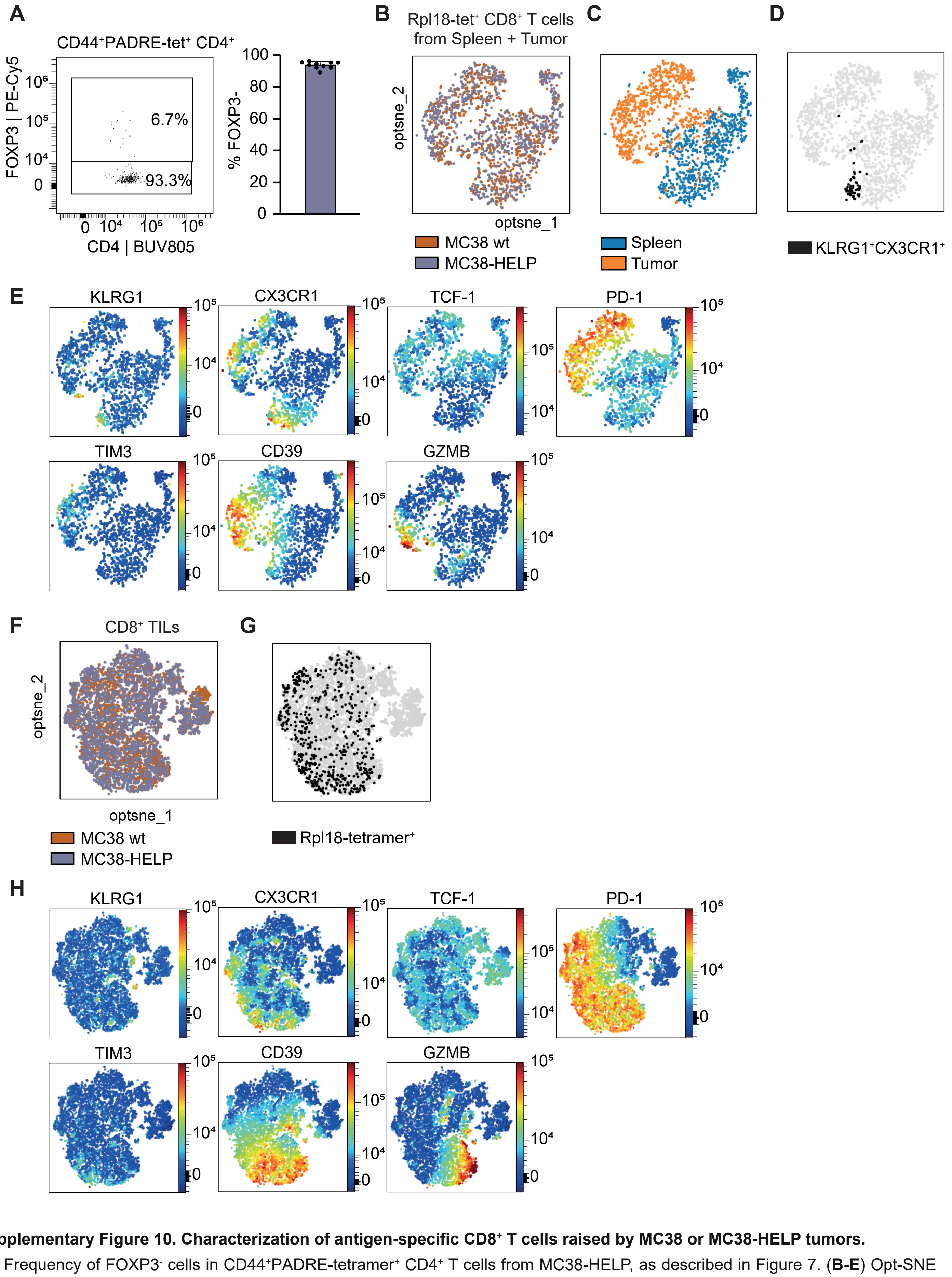


**Supplementary Figure 10. Characterization of antigen-specific CD8⁺ T cells raised by MC38 or MC38-HELP tumors.**
(**A**) Frequency of FOXP3⁻ cells in CD44⁺PADRE-tetramer⁺ CD4⁺ T cells from MC38-HELP, as described in Figure 7. (**B-E**) Opt-SNE visualization of Rpl18-tetramer⁺ CD8⁺ T cells from spleen and tumor at day 15 after MC38 or MC38-HELP tumor implantation, as described in Figure 7. (**B, C**) Distribution of cells from different tumor types (B) and tissues (C) across opt-SNE. (**D**) Visualization of the KLRG1⁺CX3CR1⁺ effector population within opt-SNE. (**E**) Expression of differentiation markers on cumulative Rpl18-tetramer⁺ CD8⁺ T cells in spleen and tumor from the experimental settings visualized by opt-SNE. (**F-H**) Opt-SNE visualization of total CD8⁺ TILs. (**F**) Distribution of cells from different tumor types across opt-SNE. (**G**) Visualization of the Rpl18-tetramer⁺ population within opt-SNE. (**H**) Expression of differentiation markers on cumulative CD8⁺ TILs visualized by opt-SNE.
