## Supplementary Methods for "Stem-like PD-1^+^TCF-1^+^ CD8^+^ T cells result from helpless priming and rely on CD4^+^ T-cell help to complete their cytotoxic effector differentiation"

**Flow cytometry staining**

Cells from blood, spleen, BM, LN and tumor were washed with FACS buffer (PBS + 2% FCS) and stained for 30 min on ice with 1:1000 LIVE/DEAD Near IR dye (Invitrogen) or 1:500 Zombie UV (Biolegend), 1:100 APC-conjugated H-2D$^b$/E7$_{49-57}$ tetramers, APC-conjugated H-2K$^b$/OVA$_{257-264}$ tetramers or APC-conjugated H-2K$^b$/KILTFDRL (mutated Rpl18) tetramers (produced in-house at LUMC) and combinations of the following monoclonal antibodies (mAbs) for surface staining: 1:100 CD45.1-BUV395 (clone A20, BD Bioscience); 1:100 CD45-BUV496 (clone 30-F11, BD Bioscience); 1:100 KLRG1-BUV563 (clone 2F1, BD Bioscience); 1:100 CD103-BUV661 (clone M290, BD Bioscience); 1:200 CD4-BUV805 (clone GK1.5, BD Bioscience); 1:800 CX3CR1-BV421 (clone SA011F11, Biolegend); 1:200 CD4-BV421 (clone GK1.5, Biolegend); 1:100 SLAMF6-Pacific Blue (clone 330-AJ, Biolegend); 1:100 CD8α-V500 (clone 53-6.7, BD Bioscience); 1:100 CD44-BV570 (clone IM7, Biolegend); 1:50 CXCR3-BV650 (clone CXCR3-173, Biolegend); 1:50 CD127-BV650 (clone A7R34, Biolegend); 1:50 CXCR5-BV711 (clone L138D7, Biolegend); 1:200 CD8α-BV750 (clone 53-6.7, BD Bioscience); 1:100 CD44-BV785 (clone IM7, Biolegend); 1:100 CD39-PE/Dazzle594 (clone Dua59, Biolegend); 1:200 TIM3-PE/Cy7 (clone RMT-23, eBioscience); 1:200 CD43-PE/Cy5 (clone 1B11, Biolegend); 1:100 CD3-PE/Fire700 (clone 17A2, Biolegend); 1:100 CD62L-PE/Fire810 (clone W18021D, Biolegend); 1:200 CD62L-AF561 (clone MEL-14, eBioscience); 1:200 KLRG1-PE/eF610 (clone 2F1, Invitrogen); 1:200 CX3CR1-PerCp/Cy5.5 (clone SA011F11, Biolegend); 1:200 PD-1-PerCP/eF710 (clone J43, eBioscience); 1:100 PD-1-APC/Fire810 (clone 29F.1A12, Biolgend). For intracellular staining, cells were fixed and permeabilized with the FOXP3/transcription factor staining buffer set (eBioscience), according to the manufacturer’s protocol. Intracellular staining was performed for 30 min on ice in permeabilization buffer, using combinations of the following mAbs: 1:100 Helios-BUV395 (clone 22F6, eBioscience); 1:100 TCF-1-AF488 (clone C63D9, Cell Signalling Technology); 1:200 GZMB-PE (clone GB-11, Sanquin); 1:100 TOX-PE (clone TXRX10, eBioscience); 1:100 GZMB-PE/CF594 (clone GB-11, BD Bioscience); 1:100 FOXP3-PE/Cy5 (clone FJK-16s, Invitrogen); 1:800 TOX-eF660 (clone TXRX10, eBioscience); 1:200 T-bet-eF660 (clone eBio4B10, eBioscience); 1:400 T-bet-AF660 (clone eBio4B10, eBioscience); 1:200 Ki67-AF700 (clone SolA15, eBioscience).

**scRNAseq analysis**

*Preparation and quality control* - Analysis of scRNAseq and TCRseq data was performed using CellRanger (10X Genomics, v6.0.1) and Seurat (v4.3.0) in R (v4.2.3). scRNAseq reads were aligned to the mouse reference genome mm10, and barcode counting was performed using ‘cellranger count’. TCRseq reads were aligned to the mouse reference genome mm10 and consensus TCR sequences were determined using ‘cellranger vdj’. From the gene expression count matrix, TCR and BCR genes were removed using the ‘grep’ function in R with the patterns “^Tr[abgd][cvdj]” and “Ig[hkl][cvdj]” respectively. After this, the dataset was analyzed in R using the standard Seurat pipeline. Seurat object was created using ‘CreateSeuratObject’, with a minimum of 200 features per cell and 3 cells expressing each gene. HTO sequencing data was added as a separate assay to the Seurat object using ‘CreateAssayObject’ and HTO counts were normalized by CLR method using ‘NormalizeData’. HTO demultiplexing was performed using ‘HTODemux’ with a positive quantile of 0.99. As a result of this, each cell in the dataset was assigned to a global HTO classification (singlet, doublet, or negative), and a HTO classification (Day-5-Help-dLN, Day-5-Help-ndLN, Day-5-No-Help-dLN, Day 5-No-Help-ndLN, Day-10-Help-dLN, Day-10-Help-ndLN, Day-10-No-Help-dLN, Day 10-No-Help-ndLN, doublet or negative). Heatmap was created of HTO demultiplexing result using ‘HTOHeatmap’ (**Supplementary Figure 6B**). Singlets were selected using ‘subset’, and gene expression data was normalized using ‘NormalizeData’. Variable features were determined using ‘FindVariableFeaures’ with the selection method ‘mean.var.plot’. Data was scaled using ‘ScaleData’ on variable features. Percentage of mitochondrial gene expression per cell was determined using ‘PercentageFeatureSet’ with the pattern “^mt-“. Singlets were filtered on number of RNA features (between 400 and 5000), number of RNA counts (<25000) and mitochondrial gene expression (<10%). Cell cycle scores for G2/M and S phase were assigned using ‘CellCycleScoring’, based on expression of G2/M and S phase genes. Data was normalized using ‘NormalizeData’ with normalization method ‘LogNormalize’, scale factor 25000. Top 2000 variable features were determined using ‘FindVariableFeatures’ with the selection method ‘vst’. Data was scaled using ‘ScaleData’, whereby G2M scores, S scores and percentage of mitochondrial gene expression were regressed out. Principle component analysis (PCA) was run using ‘RunPCA’ on variable features. Clustering was performed on first 20 principle components (PCs) using ‘FindNeighbors’ and ‘FindClusters’. Clustering resolution was set to 0.5. Uniform manifold approximation and projection (UMAP) visualization was performed using ‘RunUMAP’ on the first 20 PCs. Expression of known marker genes was visualized per cluster and within the UMAP projection using ‘Dotplot’, ‘VlnPlot’ and ‘FeaturePlot’ functions. Clusters with contaminating B cells (expression of

*Cd19, Ms4a1*), APCs (expression of *Cd40, Cd274, Cd80, Cd86*) and low quality cells (high percentage mitochondrial genes, low RNA counts) were removed. The pipeline was run again on filtered T cells, including determining variable features, data scaling, PCA, clustering and UMAP visualization. Clustering and UMAP visualization was performed on the first 50 PCs. $CD8^+$ T-cell clusters were selected by expression of *Cd8a, Cd8b1*, in absence of expression of *Cd4* (**Supplementary Figure 6C**). Within the $CD8^+$ T-cell subset, number of cells per HTO was determined (**Supplementary Figure 6D**). HTOs containing <30 $CD8^+$ T cells were excluded from further analysis.

*Gene expression analysis* - The pipeline was run again on $CD8^+$ T cells from selected HTOs, including determining variable features, data scaling, PCA, clustering and UMAP visualization. Clustering and UMAP visualization were performed on the first 20 PCs. UMAP embeddings, cluster annotations and HTO annotations per cell were exported from Seurat, and visualized using Loupe Browser (v6.4.1, 10X Genomics). Marker genes per cluster were calculated using ‘FindAllMarkers’, selecting only upregulated genes expressed in >25% of cells in the cluster, with a logFC threshold of 0.25. $CD8^+$ T-cell clusters were annotated based on expression of marker genes per cluster, as well as the expression of manually selected genes with well-known functions in T-cell differentiation according to the literature. Functional classification of differentially expressed genes between Cluster 2 and Cluster 3 was performed using Ingenuity Pathway Analysis (Qiagen).

*TCRseq analysis* - Sequences of CDR3 and VDJ gene annotations of paired TCRα and TCRβ chains were analyzed using Loupe VDJ Browser (v5.0.0, 10X Genomics). Expanded clones were identified as TCR clonotypes containing 2 or more cells. TCR sequences from Loupe VDJ Browser, and UMAP projection and cluster annotation from Seurat were visualized together using Loupe Browser (v6.4.1, 10X Genomics). For determining expanded clonotypes per differentiation cluster, only cells that grouped together in the UMAP with their assigned cluster were included for quantification.

*Trajectory analysis* was performed using Seurat and Monocle3 (v1.3.1) in R. Seurat object of $CD8^+$ T cells from selected HTOs was converted into cell data set object using ‘as.cell_data_set’, after which partitions, cluster annotations and UMAP embeddings were added manually. Trajectory was plotted using ‘learn_graph’, after which the trajectory branch of interest (from Cluster 0 to Cluster 3) was selected using ‘choose_graph_segments’. Pseudotime was calculated using ‘order_cells’, and pseudotime values per cell were exported for visualization using GraphPad Prism and Loupe Browser.

For *comparison of our scRNAseq data to published gene signatures*, top 200 marker genes of TCF1$^+$ progenitor (cluster 9) and effector (cluster 6) CD8$^+$ T cells from chronic and acute LCMV infection were downloaded from Table S5 from Pritykin *et al*. 2021[25]. These top 200 genes were used to calculate a module score for each cell in our CD8$^+$ T cells from selected HTOs, using the function 'AddModuleScore' in R. The same method was performed using top 100 marker genes of stem-like CD8$^+$ T cells from the TdLN, (cluster 4) as published by Connolly *et al.* 2021[67], which were provided upon request.
